# Origin of subgenomes in the circumboreal allopolyploid carnivorous plant *Drosera anglica* (Droseraceae)

**DOI:** 10.1101/2025.01.09.632192

**Authors:** Rebekah A. Mohn, Ya Yang

**Author notes:** Correspondence: Rebekah A. Mohn.

## Abstract

**Premise of Study:** The parentage of a widespread member of the carnivorous sundew genus *Drosera*, the allopolyploid *Drosera anglica,* remains uncertain despite over 100 years of morphological, cytological, and, more recently, molecular study.

**Methods:** Using transcriptomic and genomic data from 12 species *Drosera* sect. *Drosera* including four *D. anglica* populations and a disjunct Idaho population of *D. intermedia*, we assembled genes in HybPiper and phased sequences in HybPhaser. We estimated heterozygosity and generated flow cytometry data to assess ploidy levels. We estimated species relationships with phylogenetic and pairwise genetic distance methods. Additionally, we assembled *rbc*L and ITS reads to compare to previous data.

**Key Results:** Sequences from phased subgenomes highly supported *D. anglica* as sister to *D. rotundifolia* and *D. linearis,* differing from previous analyses based on chromosome pairing and Sanger sequencing with limited taxon sampling. Both ITS and *rbc*L sequences of *D. anglica* were the most similar to *D. linearis*. *Drosera anglica* is intermediate between both parents in leaf shape and microhabitat; however, across *D.* sect. *Drosera*, neither leaf shape nor biogeographic distribution were reliable indicators of phylogenetic relationships. Despite a range-wide sampling, we did not find evidence for multiple origins of *D. anglica*. Additionally, we confirmed that the Idaho population previously identified as *D. intermedia* is *D. anglica*.

**Conclusions:** *Drosera anglica* arose from allopolyploidy between *D. linearis* (the chloroplast donor) and *D. rotundifolia*. Our study demonstrates the importance of taxon sampling and careful examining complex phylogenomic data, and presents an exemplar of analyzing allopolyploid relationships in plant lineages.

## INTRODUCTION

Allopolyploidy is a driving force of evolutionary innovation. It brings together genes, regulatory elements, and variants in ways that differ from either diploid parent (Nieto Feliner et al., 2020). As the allopolyploid genome undergoes subsequent restructuring, gene copies may be lost, exposed to new regulatory elements, or located in different epigenetic contexts (Nieto Feliner et al., 2020). To understand the long-term trajectory of these changes and their impact on species diversification, we need to identify the parental species and lineages. A combination of careful morphological, cytological, and sequence approaches, complemented by distribution and habitat information, is ideal, as each of these lines of evidence on its own can be insufficient or even misleading.

*Drosera* L. (Droseraceae, Caryophyllales) is a carnivorous plant genus of ∼250 species worldwide. Twenty-eight of those species are in *D.* subg*. Drosera* sect. *Drosera* based on molecular, cytological, and morphological evidence (Fleischmann et al., 2018). This section is the most geographically widespread of the genus: found in Europe, Asia into Oceania, North America, and South America (Fleischmann et al., 2018; Lowrie et al., 2017). One of these species alone, *D. anglica* or the English sundew, is found in boreal North America, Europe, Asia, and Hawaii (Fleischmann et al., 2018; Lowrie et al., 2017). All eight species of *Drosera* native to North America (Mellichamp, 2016) belong to *D.* sect. *Drosera* (Figure 1). These eight species are all carnivorous and found in nitrogen and phosphorous poor habitats with two general geographic distributions. Five species have a primary north-south distribution along the East Coast of North America, with three of these extending into South America (Mellichamp, 2016).

**Figure 1:**
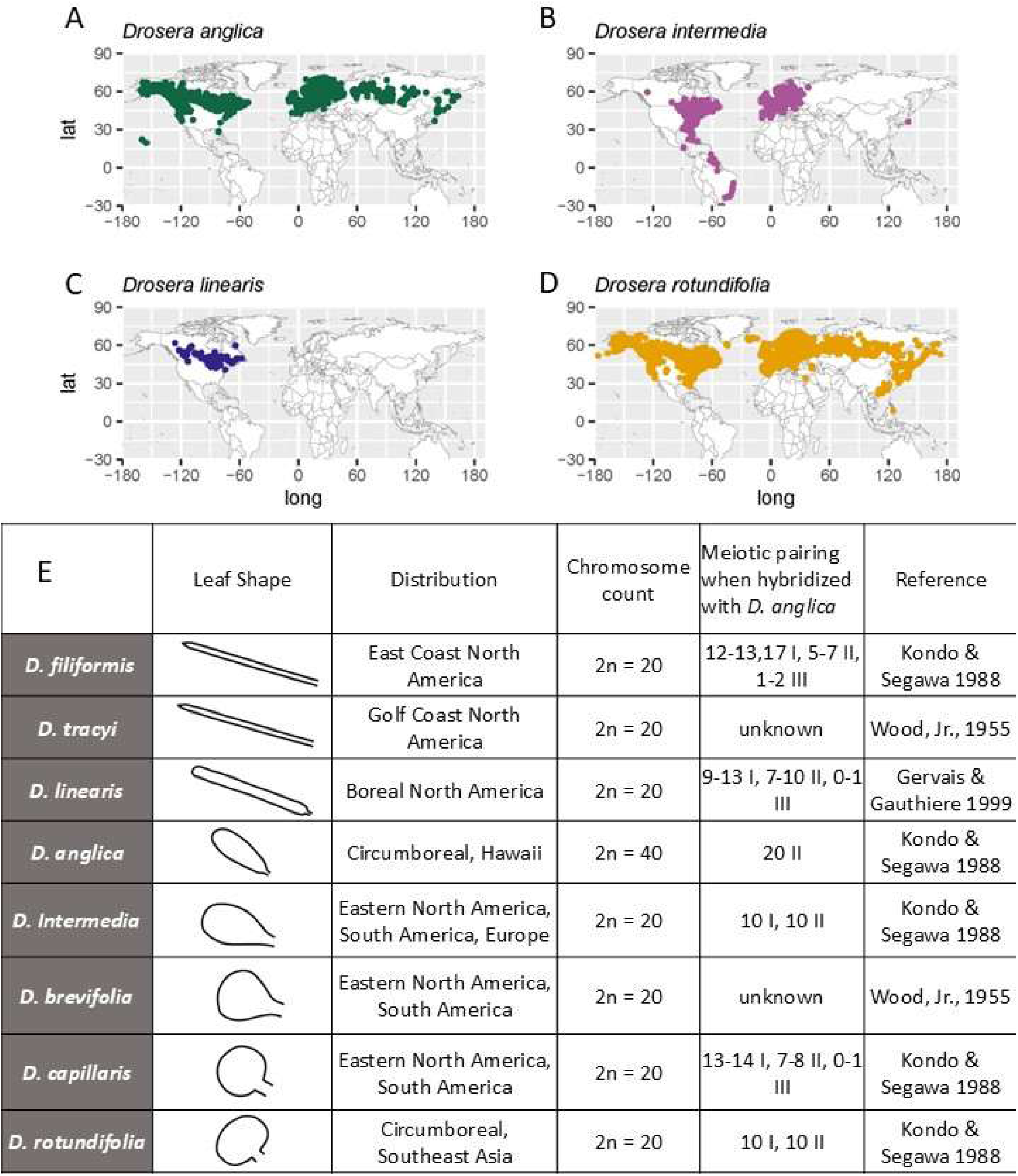
Distribution and cytogenetics of North American *Drosera.* A-D) Distribution of cleaned GBIF records (*GBIF Occurrence Download*, 2024) of *D. anglica* and putative parental species, *D. rotundifolia*, *D. linearis* and *D. intermedia* (the Idaho population not shown). E) A table summarizing the leaf shape, distribution (Lowrie et al., 2017; Mellichamp, 2016), and cytological data of the eight North American *Drosera* species. Chromosome pairing of meiotic counts in hybrids between *D. anglica* and five diploid species are listed as the number of univalent (I), bivalent (II), and trivalent (III) chromosome pairs.

Among them, *Drosera intermedia* is also found in Europe. The remaining three species have east-west distributions in the boreal zone: both *D. anglica* and *D. rotundifolia* are circumboreal with disjunct populations in tropical mountains, and the third species, *Drosera linearis*, is restricted to boreal North America (Figure 1).

North American *D.* sect. *Drosera* vary in leaf shape and orientation. Leaf blades range from long and narrow in *D. filiformis* and *D. linearis,* spatulate in *D. intermedia,* to short and round or wider than round in *D. capillaris, D. brevifolia,* and *D. rotundifolia* (Figure 1). Leaves of *D. brevifolia, D. capillaris,* and often *D. rotundifolia* are appressed to the ground, while leaves of *D. filiformis, D. linearis, D. anglica,* and *D. intermedia* are raised to erect. Leaf shape and orientation are thought to be associated with habitat and prey (Fleischmann et al., 2018); for example, raised leaves with long petioles can help the lamina stay above the water when water levels rise.

All North American *Drosera* species have a diploid chromosome number of *2n* = 20, except *D. anglica*, which has a tetraploid chromosome count of *2n* = 40. In 1903 with the burgeoning study of chromosomes in hybrids, Rosenberg (1903, 1904, 1909) noticed that during meiosis in hybrids between *D. rotundifolia* (2*n* = 20) and *D. anglica* (2*n* = 40), there were 10 bivalent (II) and 10 univalent (I) chromosome pairs (from this point referred to as 10II, 10I). The 10 II chromosomes are likely properly paired homologous chromosomes, while the 10 I chromosomes are unpaired. This observation led to the hypothesis that *D. anglica* was a hybrid polyploid between *D. rotundifolia* and another species (Winge, 1917). This raised the question of the identity of the other parental species and led to subsequent cytological work on hybrids between *D. anglica* and most of the other North American *Drosera* species (Figure 1; Gervais & Gauthier, 1999; Kondo & Segawa, 1988).

Of the candidate species, *D. intermedia* and *D. linearis* were most likely to be the second parental lineages given their overlapping distribution and similarity in habitat and morphology compared to *D. anglica* (Figure 1). *Drosera anglica* and *D. intermedia* are easily confused, especially in herbarium specimens, as their leaf shapes are very similar, and the plants, especially early in the vegetative stage, have few distinguishing features. *Drosera intermedia* occurs in eastern North America, Europe, and South America. It has been reported from two disjunct populations in northern Idaho and southern Idaho more than 1000 km from the nearest populations (Lowrie et al., 2017; Mellichamp, 2016). The Idaho populations have been protected by the state as a rare plant with potentially unique genetics, but some botanists have suggested that they may be misidentified *D. anglica* populations (Lowrie et al., 2017; Rice, 2019). While *D. rotundifolia* and *D. intermedia* rarely form hybrids with each other in the wild (Grima, 2020), *D. anglica* and *D. intermedia* form 10 II and 10 I pairs when hybridized artificially (Kondo & Segawa, 1988). Work comparing isozymes of European *Drosera* found that *D. anglica* and *D. rotundifolia* shared allozymes and *D. intermedia* and *D. anglica* did not, contradicting the evidence from morphology and chromosome pairing (Seeholzer, 1993).

Alternatively, *D. linearis* has been hypothesized as the other parent of *D. anglica* based partly on the similarity of their microhabitats. *Drosera anglica* occurs in fen strings (ridges in patterned fens), fen edges, and wetter regions of bogs. This microhabitat is intermediate between the often calcium-rich fen flark (a depression in a patterned fen) microhabitat of *D. linearis* and the drier, more acidic sphagnum hummock microhabitat of *D. rotundifolia. Drosera anglica* is also intermediate in leaf shape between *D. linearis* and *D. rotundifolia* (Figure 1; Wood, Jr., 1955). However, the chromosomes of *D. linearis* do not pair properly with those of *D. anglica* (Figure 1). When hybridized in the wild, *D. anglica* ✕ *D. linearis* forms 9-13 I, 7-10 II, and 1 III during meiosis (Gervais & Gauthier, 1999). Chromosomes in hybrids between *D. rotundifolia* and *D. linearis* do not pair properly either (2-10 I, 3-7 II, and 1-3 III; Wood, Jr., 1955). Although fertile neo-allopolyploid hybrids of *D. rotundifolia* and *D. linearis* have been found in nature, they are slightly different in appearance compared to *D. anglica* (Wood, Jr., 1955). Additional work on the flower structure of *D. anglica* found that it was not always intermediate between *D. rotundifolia* and *D. linearis* with, for example, *D. anglica* having the smallest and fewest papillate cells on the anther apices of the three species (Murza & Davis, 2003).

Despite a long history and large body of cytological work, previous molecular phylogenetic work on *Drosera* has been inadequate to clarify *D. anglica*’s parentage. Rivadavia et al. (2003) found that the *rbc*L sequence of *D. anglica* and *D. rotundifolia* only differed in three base pairs, supporting the hypothesis that *D. rotundifolia* was the maternal parent. However, this and other more recent phylogenetic studies of *Drosera* have all relied on two to three loci and lacked phylogenetic signal for resolving the relationships within the recently diversified *D.* sect. *Drosera* (Rivadavia et al., 2003; Veleba et al., 2017). The absence of *D. linearis* in previous phylogenetic or allozyme studies due to its restricted distribution and remote habitat has also left the parentage of *D. anglica* unresolved.

In order to detect both maternal and paternal parents of *D. anglica* in the context of the recently diverged *D.* sect. *Drosera,* increased sampling of nuclear loci is needed. Transcriptomes provide thousands of loci each with relatively long sequences informative for evaluating discordance among taxa with short branch lengths (Yang et al., 2015). It is also an effective genome subsampling approach to tease apart subgenomes. Additionally, transcriptome datasets can be re-analyzed and combined with genome resequencing and target enrichment datasets or used later to address molecular evolution questions.

With this information in mind, we sequenced transcriptomes from one to four populations each of 12 species from across *D*. sect. *Drosera*. We reconstructed gene trees and species trees, phased subgenomes, and analyzed genetic distance and allele diversity to answer the following questions: 1) What are the maternal and paternal species or lineages of *Drosera anglica,* and how does that compare to previous cytological findings? 2) Is the northern Idaho population of ‘*D. intermedia’ D. intermedia* or *D. anglica*? and 3) Since *Drosera anglica* is circumboreal and occurs in Hawaii, did it originate once and spread subsequently through long-distance dispersal or is there any evidence for multiple origins?

## METHODS

### Sampling and Collection

To maximize the chance that our sampling included the parents of *D. anglica*, we sampled species from across *D.* sect. *Drosera*, with an emphasis on potential parents of *D. anglica.* We selected species based on the previously published *rbc*L phylogeny (Rivadavia et al., 2003) representing major clades, different geographic distributions, and a diversity in leaf morphology. For *D. anglica,* we sampled across its distribution and included multiple populations from its hypothesized parents.

We sampled one European, one Hawaiian (Morden et al., 1996; Randell & Morden, 1999), and two North American populations of *D. anglica*. We also sampled the northern Idaho population, whose identification has been disputed in the recent literature and will here refer to it as ‘*D. intermedia’.* Additionally, we sampled two populations each of *D. linearis*, *D. rotundifolia* and *D. capillaris,* and one population for three of the four remaining North American species and four South American species. To round out our dataset, we downloaded a publicly available transcriptome from a Russian population of *D. rotundifolia* (NCBI SRA: SRR8948654; Gruzdev et al., 2019). In total, we sampled seven out of the eight North American species. The eighth species, *D. tracyi*, is morphologically similar to and has at times been considered a subspecies of *D. filiformis* with a more restricted range than *D. filiformis* (Lowrie et al., 2017; Rice, 2011). For clarity, we will refer to each sample by the species name, and for species with multiple collections, the species name is followed by the location abbreviation in parentheses. For outgroups, we used the published *D. spatulata* reference genome (Palfalvi et al., 2020) since it is diploid and sister to the rest of *D.* sect. *Drosera* (Veleba et al., 2017); we also used *D. prolifera*, a diploid in *D.* subg. *Drosera* but outside of *D.* sect. *Drosera* (Fleischmann et al., 2018; Veleba et al., 2017).

Newly generated transcriptomes came from field-collected and cultivated samples. For most North American samples, we collected the whole plant with roots and some substrate in the field and placed it in a Ziploc bag. When back at the trailhead, we transferred the above-ground portion of the plant into an 8 mL Nalgene bottle and froze the sealed bottle in liquid nitrogen.

Depending on the hiking distance, samples were frozen within 3 hours of collection. Cultivated plants were sourced from a reputable specialty grower (Best Carnivorous Plants; www.bestcarnivorousplants.net) for South American and European populations (Table 1). Tissue from cultivated plants was collected, immediately placed in a 2 mL lysing tube with Lysing Matrix A (MP Biomedicals), and flash frozen in liquid nitrogen. To avoid contamination while collecting the samples, we wore gloves, and between species cleaned tweezers with 70% ethanol followed by RNase*Zap* from Invitrogen by Thermo Fisher Scientific. If our gloves came in contact with the plant, we also changed our gloves. All samples were ground using the FastPrep-24™ 5G bead beating grinder (MP Biomedicals) and Lysing Matrix A (MP Biomedicals) in dry ice with the CoolPrep™ adapter (MP Biomedicals). RNA was extracted using the PureLink™ Plant RNA Reagent (Invitrogen; see Appendix S1 for modifications to the manufacturer’s protocols; Yang et al., 2017). Library preparation and sequencing was done by either the University of Minnesota Genomics Center (UMGC) or Novogene Corporation, Inc. (Appendix S2). At the UMGC, the libraries were prepared using Illumina Ribo-Zero Plus rRNA Depletion Kit and were sequenced either with 125 paired-end reads on the Illumina HiSeq 2500 or with 150 single-end reads on the NextSeq 550. We requested paired-end reads, but the sequencing facility accidentally performed single-end read library prep, and the remaining samples were discarded. Novogene used the New England Biological NEBNext Ultra II Directional RNA Library Prep kit and sequenced 150 paired-end reads on the Illumina NovaSeq 6000 platform.

**Table 1:**
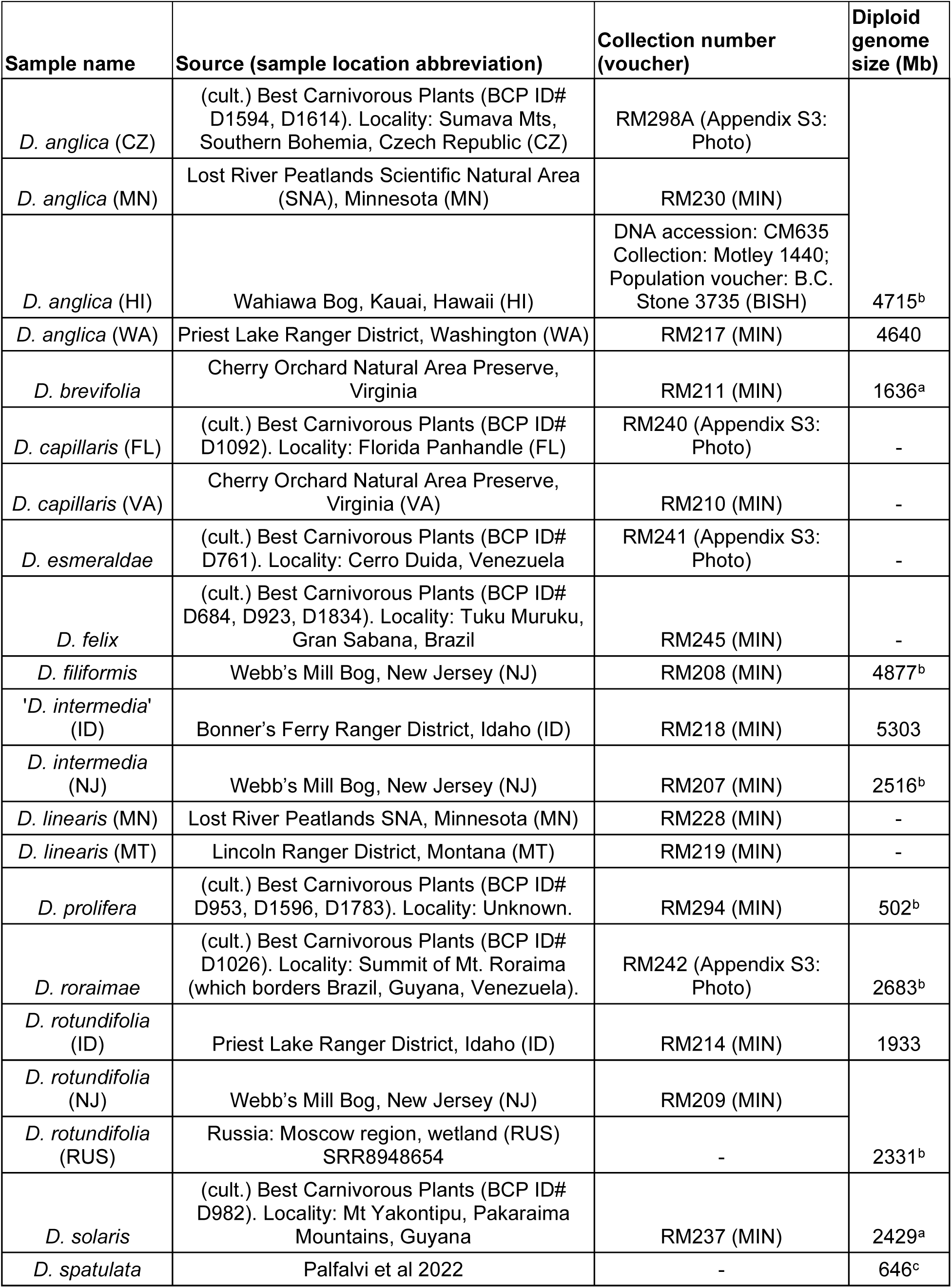
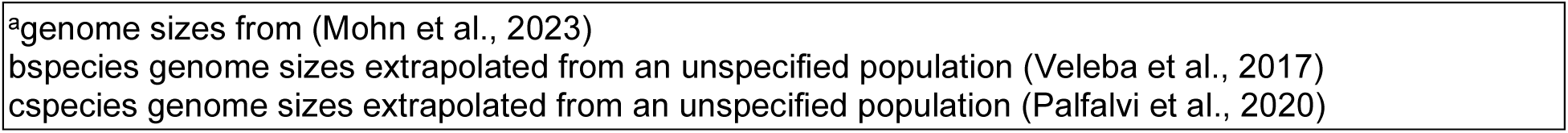
Sample source, voucher information, and genome sizes. The collection locality is in the USA when unspecified.

For a Hawaiian population of Drosera anglica, we obtained DNA from the Hawaiian Plant DNA Library (Morden et al., 1996; Randell & Morden, 1999). We sequenced this sample at Novogene with AB clonal DNA Library Prep Kit, 350 bp size selection after fragmentation and 150 paired-end reads on the Illumina NovaSeq 6000 platform.

### Observation data

We downloaded “Human observation” and “Preserved specimen” GBIF data with an occurrence status of “present”, and either a “Coordinate rounded” or “WGS1984 inferred” flag for *D. anglica, D. intermedia, D. linearis,* and *D. rotundifolia* (*GBIF Occurrence Download*, 2024). After downloading, observations that had a “Country coordinate mismatch” were removed. Map trimming also removed a single observation from New Zealand that is likely due to misidentification or an escape from cultivation.

### Genome size estimation

Fresh samples were collected and mailed to the Flow Cytometry Core Lab at the Benaroya Research Institute (Seattle, WA, U.S.A.) for genome size estimation. Samples were stained with propidium iodide (PI) and analyzed with a FACScalibur flow cytometer (Becton-Dickinson, San Jose, CA) following the protocol described by (Arumuganathan & Earle, 1991)). For each sample, four flow cytometry measurements were taken with an internal size standard (see Table S1). We used the average genome size for each sample for subsequent analyses. The source, voucher, and size standards used for generating new flow cytometry data are listed in Table S1.

### Read processing and sequence assembly

For the transcriptomic data, except where noted, we followed the previously published pipeline https://bitbucket.org/yanglab/phylogenomic_dataset_construction/ (Morales-Briones et al., 2021; Yang & Smith, 2014) to clean reads, assemble transcripts, and carry out phylogenomic analysis. Programs Rcorrector (Song & Florea, 2015), Trimmomatic (Bolger et al., 2014), Bowtie2 (Langmead & Salzberg, 2012), and FastQC (Andrew, 2010) were used to clean, trim, map, and filter out organellar reads, and detect and filter over-represented reads.

For the genomic data, we filtered organellar reads. To reduce the data to only relevant reads, we used BBmap allowing 10 substitutions per read and 80% identity to map reads to consensus assemblies of *D. rotundifolia* (NJ) and *D. linearis* (MN) generated by HybPhaser.

### Distribution of synonymous distance (Ks) estimated using de novo assembled transcriptomes

The cleaned transcriptome reads were *de novo* assembled with Trinity version 2.5.1 (Haas et al., 2013). To test for cross-contamination, the cleaned reads and the assembled transcripts were fed into Croco version 1.1 (Simion et al., 2018) in two groups: one from paired-end and the other from single-end datasets as Croco can only take one type of read configuration in each run.

To estimate the timing of polyploidy events with Ks plots, assembled transcripts from Trinity were translated with TransDecoder version 5.3.0 (Haas, BJ, 2015/Accessed: 2018; https://github.com/TransDecoder/TransDecoder) without any filtering. Within-species Ks plots were calculated following (Yang et al., 2015, 2018). We removed Ks values < 0.01 before visualizing Ks distributions as heterozygosity and isoforms often contribute to Ks values less than 0.01.

Visual inspection of reads mapped to initial assemblies indicated that the reads from the single-end sequencing batch (*D. rotundifolia* (ID), *D. anglica* (WA), and ‘*D. intermedia*’ (ID)) often had a T at the 3’ end, likely as part of the adapter sequence. Therefore, one base was removed from the 3’ end of these three samples for subsequent analyses unless otherwise stated.

### Synthetic in silico hybrids

As a positive control to evaluate our ability to tease apart subgenomes in allopolyploids, after cleaning the raw reads, we combined 6,666,667 read pairs from *D. linearis* (MT) and 8,000,000 read pairs from *D. rotundifolia* (NJ) to make a synthetic *in-silico* hybrid that roughly resembled *D. anglica* in read coverage. This difference in the number of reads made up for the 125 PE reads of *D. rotundifolia* (NJ) and 150 PE reads of *D. linearis* (MT). Since single-end reads may suffer from additional challenges in phasing, we made a second synthetic hybrid with the same reads but without pairing information. From now on, these two datasets will be referred to as synthetic hybrids.

### Selecting target genes for targeted assembly using HybPiper

We chose targeted assembly for phylogenetic analyses given the tools available for phasing subgenomes, and the ability to combine our genome and transcriptome datasets. To choose target genes that are single-copy and well supported by transcriptome data, and to reduce computational time, an initial round of phylogenomic analysis was performed with transcriptome assemblies from two *D. linearis*, two *D. rotundifolia* (before the 3’ end of *D. rotundifolia* (ID) was trimmed), and *D. intermedia* (NJ) and the CDS file from the *D. spatulata* genome annotation (Palfalvi et al., 2020). This was done following the Yang & Smith pipeline (Morales-Briones et al., 2021; Yang & Smith, 2014).

We used TransRate version 1.0.3 (Smith-Unna et al., 2016) to quantify the quality of the Trinity assemblies and removed transcripts with nucleotides of mapped reads poorly matching the assembled transcript (s(Cnuc) ≤ 0.25), low read coverage (s(Ccov) ≤ 0.25), or paired-end reads misaligned (s(Cord) ≤ 0.5) in paired-end read samples. For all chimeric transcripts with multiple open reading frames stitched together in opposite directions, each with at least 30% similarity in at least 100 bp of the CDS compared to *Beta vulgaris* proteins were removed (Yang & Smith, 2013). The resulting transcripts were translated with TransDecoder with *Arabidopsis thaliana* and *Beta vulgaris* reference proteomes. Finally, the CDS were further reduced with CD-HIT-EST (W. Li & Godzik, 2006) to remove sequences with > 99% similarity using a 10-bp word length.

To cluster the resulting CDS, hits from an all-by-all BLASTn (Altschul et al., 1990; Camacho et al., 2009) search with at least a 30% length coverage of both sequences were input into mcl (Van Dongen, 2008) with an inflation value of 1.4. The resulting clusters were each aligned with MAFFT version 7.475 (Katoh & Standley, 2013) using the generalized affine gap cost for pairwise alignments with 1000 iterations, alignments trimmed with Phyx (Brown et al., 2017) removing columns with >90% missing data, and gene trees estimated using RAxML (version 8.2.11). Using TreeShrink (Mai & Mirarab, 2018) we trimmed spurious tips that were in the 0.4 quantile, and then we removed monophyletic tips of the same sample, leaving only one with the highest number of aligned characters in the trimmed alignment. We then visually inspected the resulting gene trees and cut long internal branches that were longer than 0.1 substitutions per site, as internal branches among our sampled species were mostly < 0.06, and branches longer than 0.1 were due to deeper paralogs or spurious sequences. We retained trees with all six taxa, and re-aligned the sequences with MAFFT. Terminal branches 10× longer than their sister or more than 1.0 substitutions per site were trimmed. We thinned monophyletic and paraphyletic tips from the same sample to the one with the highest number of aligned characters in the trimmed alignment. Finally, we selected one-to-one orthologs present in all six samples and used the *D. spatulata* coding sequences for these genes in downstream analyses.

### Targeted assembly with HybPiper

As part of the HybPiper 2.0 (Johnson et al., 2016) pipeline with default settings, we used bwa (H. Li, 2013) to map reads to the 6443 *D. spatulata* target genes selected as above, and SPAdes (Bankevich et al., 2012) to assemble the mapped reads into transcripts, requiring a read depth ≥ 4 given the uneven read coverage of transcriptome data. Assembled transcripts for each gene were compared to each other and the reference using Exonerate version 2.2.0 (Slater et al. (http://www.ebi.ac.uk/~guy/exonerate/; Slater & Birney, 2005) and were retained if the hits had ≥ 85% identity. If multiple transcripts were assembled by SPAdes that each covered >50% of the length of the target, HybPiper throws a long paralog warning. For genes without an assembled (“long”) paralog but with a second contig that covered <50% of the target length and with a read depth of 1–4 for the unassembled length of the gene, HybPiper throws a depth paralog warning.

### Reference-based phasing with HybPhaser

We used the HybPhaser pipeline to identify polyploids, identify parental lineages, and phase the subgenomes (Nauheimer et al., 2021). This consists of four steps: visualizing the distribution of single nucleotide polymorphisms in each sample to detect potential hybrids and polyploids, determining whether samples belong to multiple clades by mapping to clade references, phasing reads to the appropriate clade references, and then re-assembling the phased reads in HybPiper. First, loci with less than 20% of samples or covered <10% of the length of the target sequence were removed. Using only the reads mapped to the target by bwa in HybPiper, HybPhaser version 2.1 (Nauheimer et al., 2021) generated a consensus sequence for each gene in each sample. For a second allele to be called at a site according to the default settings, there must be a read depth of at least 10 at the site and the second allele must be supported by at least four reads and 15% of the reads. Allele divergence, the percentage of SNPs per gene length, and loci heterozygosity, the percentage of loci with SNPs, were calculated by HybPhaser. In single-copy genes in diploid species, allele divergence equals the nucleotide diversity per site (π). In genes with multiple closely related copies, especially in polyploid species, paralogs and homeologs also contribute to allele divergence. We identified samples with elevated heterozygosity and sequence divergence as evidence of polyploidy and/or hybrids and candidates for subgenome phasing. To select which dataset to use as clade references to phase to, we gathered the genes from the diploid individuals of the initial HybPiper run, aligned with MAFFT v7.475 (Katoh & Standley, 2013), trimmed alignment with Phyx (Brown et al., 2017) removing columns with >90% missing data, estimated gene trees in RAxML version 8.2.11 (Stamatakis, 2014), and estimated the species tree in ASTRAL version 5.7.8 (Zhang et al., 2017). We selected references that had low heterozygosity and low sequence divergence and represented different clades within *D.* sect. *Drosera*. Every sample was then mapped to the consensus assemblies of the clade references. Samples that had both high heterozygosity and sequence divergence and mapped to multiple clade references at approximately equal rates were then phased. To phase subgenomes, HybPhaser mapped reads from a sample to both references consensus assemblies using BBMap version 38.96 (Bushnell, 2014). If the reads are mapped unambiguously to one of the references, the reads are sorted to that reference. If they map to both equally, they go to both. If they did not map to either, they were removed. This produced two files with phased reads that were each assembled to the original 6443 *D. spatulata* genes using HybPiper.

The resulting phased homeologs from HybPhaser and unphased genes from the remaining samples were again aligned using MACSE_OMM v. 12.01, which takes codon sequence into account (Ranwez et al., 2011, 2021), and alignments trimmed using Phyx (Brown et al., 2017) requiring a minimum of 5% column occupancy. A subset of resulting alignments were visually inspected to ensure proper assembly and phasing. RAxML was used to estimate gene trees with 100 bootstrap replicates. ASTRAL was then used to estimate the species tree from the gene trees. Gene tree discordance was then calculated by PhyParts (Smith et al., 2015) requiring a minimum local bootstrap of 50. Trimmed alignments with at least 50 bp in aligned length were concatenated for phylogenetic reconstruction using RAxML.

### Pairwise genetic distance

In addition to tree-based methods, we also calculated the pairwise genetic distance between samples. A distance-based method is informative, especially when relationships among samples are not strictly tree-like and when samples are very closely related. We removed the MACSE-OMM alignments with less than 98.0% of pairwise identical base pairs in the alignment (including gap to base pair comparison), which removed alignments with large segments of ambiguous characters or high levels of missing data. After visually inspecting the remaining alignments, we removed alignments with ambiguous characters, removed aligned columns with any gap, and kept only alignments longer than 1000 bps to ensure enough signal. We then used bio3d version 2.4-4 (Grant et al., 2006) in R version 4.2.3 to calculate the pairwise genetic distance of each sample or subgenome per alignment. We then calculated the mean and median genetic distance between each pair of samples or subgenomes.

### *rbc*L and rRNA Sequence Variation

To compare our results with previously published phylogenetic studies based on Sanger sequencing, we extracted the chloroplast *rbc*L and nuclear rRNA raw reads from our *D. linearis*, *D. rotundifolia*, *D. anglica,* and ‘*D. intermedia*’ transcriptomes. To ensure that SNPs were not due to low-quality bases, we first performed a second, more stringent round of Trimmomatic trimming with settings LEADING:28 TRAILING:28 SLIDINGWINDOW:8:28 SLIDINGWINDOW:1:10 MINLEN:65. For rRNA, we used Bowtie2 to map the trimmed reads to a *D. rotundifolia* sequence MT784099.1 from NCBI GenBank. For *rbc*L, we used the extracted organellar reads (pre 3’ end trimming and second Trimmomatic trimming) and Bowtie2 to map the reads to a reference *rbc*L sequence from *D. rotundifolia* (AB29809.1 from NCBI GenBank). In both cases we used the Bowtie2--end-to-end setting for all samples and --no-mixed setting for paired-end samples. We called variants using BCFtools mpileup with default settings, except that the max depth was increased to 30,000 and visualized the mapping and variants in the Integrative Genomics Viewer version 2.12.3 (IGV).

## RESULTS

### Sampling, sequencing, assembly, and initial quality check

Of the 28 species in *D.* section *Drosera* (Fleischmann et al., 2018), we sampled transcriptomes or genomes from 12 species in 20 populations. Of these, 17 were newly generated transcriptome datasets, and one was a newly generated genome sequencing dataset. The RNA integrity number (RIN) from the successfully sequenced RNA samples ranged from 2.0 to 7.3.

For each dataset, raw sequencing reads ranged from 22 million to 26 million single-end reads or 20 to 44 million paired-end reads (Table S1). Initial, unphased targeted assembly with HybPiper recovered >4737 genes from ingroup samples with paired-end reads. After reference-based phasing with HybPhaser, the second round of HybPiper assembly recovered >5834 genes for each subgenome from both single-end and paired-end in-group samples.

Croco found no evidence of cross-contamination among paired-end samples. For the single-end samples, Croco flagged 2–13% of assembled transcripts from three samples with potential cross-contamination. These three samples (*D. rotundifolia* (ID), *D. anglica* (WA), and *’D. intermedia’* (ID)) are too genetically similar to distinguish cross-contamination from identical short reads. Therefore, we proceeded with the analysis using all the samples.

All transcriptomes shared a Ks peak at around 1.8 and a second, more recent Ks peak at around 0.7, corresponding to the shared whole genome duplication events early in Eudicots and early in Droseraceae, respectively (Palfalvi et al., 2020; Walker et al., 2017). No additional, more recent Ks peaks were apparent in any of the samples (Appendix S4).

### Few *D. anglica* paralogs were detected by HybPiper despite high nucleotide diversity

Gene clustering and tree-based ortholog inference using *D. spatulata* CDS from genome annotation and *de novo* assembled transcripts from *D. intermedia, D. rotundifolia,* and *D. linearis* resulted in 6443 genes with one copy per sample to be used as targets for HybPiper assembly. Of the 6443 target genes, 4497 to 6393 were assembled in each sample. Of these, HybPiper returned paralog warnings for 92–841 genes on multiple long assembled transcripts and 112–2311 genes for paralogs with insufficient read coverage (< 4) to assemble to at least 50% the length of the reference (unassembled from here on). Paired-end *D. anglica* samples and the paired-end synthetic hybrid had a higher number of assembled (501–841 versus < 229) and unassembled (1805–2311 versus < 500) paralogs compared to other paired-end samples. In the *D. anglica* sample with single-end reads, only 320–332 assembled and 456–458 unassembled paralogs were detected. Both synthetic hybrids had similar numbers of paralogs to *D. anglica* samples (493–557 assembled; 848–921 unassembled), but not as high as might be expected for a polyploid indicating that homeologs were not properly separated in allopolyploids.

Visualizing SNPs as the initial steps of the HybPhaser pipeline found that all *Drosera anglica* samples, ‘*D. intermedia*’, and the synthetic hybrids had much higher loci heterozygosity and allele divergence than diploid samples of *D.* sect. *Drosera*. Loci heterozygosity, the percentage of genes with SNPs, ranged from 97% to 99% in *Drosera anglica* and ‘*D. intermedia*’ (ID) while in all other samples ranged from 14% to 30% (Figure 2). Similarly, allele divergence, the percentage of SNPs per gene length, was 1.9–2.6 for *D. anglica* and ‘*D. intermedia*’ (ID), compared to < 0.6 for all other samples (Figure 2). The synthetic hybrids had both loci heterozygosity and allele divergence very similar to, albeit slightly lower than *D. anglica*. Given the highly elevated nucleotide diversity in *D. anglica* and *’D. intermedia’* (ID) samples, the relatively few paralogs detected, especially in the synthetic hybrids and single-end read samples, suggested that paralogs were too similar to be distinguished and were assembled into chimeric sequences. Subsequently, we used a reference-based phasing approach in HybPhaser to distinguish between subgenomes.

**Figure 2:**
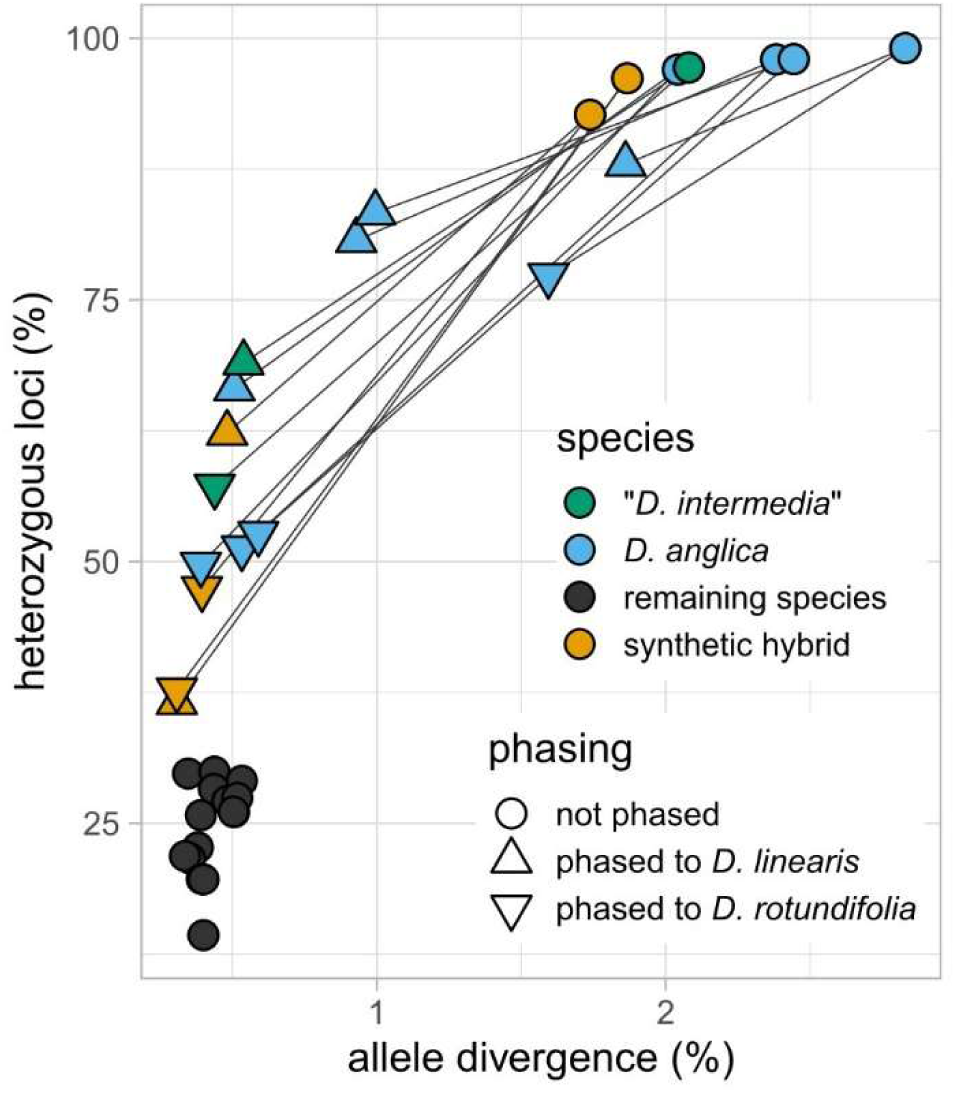
*Drosera anglica* and the *’D. intermedia*’ (ID) samples had much higher allele divergence and loci heterozygosity than the remaining species in *D.* sect. *Drosera*, at levels similar to the synthetic hybrid. After phasing the subgenome’s, each sample’s allele divergence and loci heterozygosity decreased but remained higher than the diploid species. Lines connect phased vs. unphased datasets from the same sample.

### Reference-based phasing showed *D. anglica* and ‘*D. intermedia*’ (ID) subgenomes being most similar to *D. rotundifolia* and *D. linearis*

Based on initial species tree inference, we chose *D. rotundifolia* (NJ)*, D. linearis* (MN)*, D. solaris, D. brevifolia,* and *D. filiformis* as clade references for HybPhaser (Figure 3). Most samples mapped strongly to one reference or weakly to multiple references except all samples of *D. anglica* and ‘*D. intermedia*’ (ID), which mapped strongly to both *D. rotundifolia* (NJ) and *D. linearis* (MN; Table S1.3). After phasing *D. anglica* and ‘*D. intermedia*’ (ID) reads against *D. rotundifolia* (NJ) and *D. linearis* (MN), slightly more genes were recovered for the *D. linearis* subgenome in all samples (Table S1.4). In total 3057 genes were assembled in all subgenomes and diploid samples with HybPiper. Allele divergence in the phased *D. anglica* and ‘*D. intermedia*’ (ID) samples decreased but remained higher in the *D. linearis* subgenome than the *D. rotundifolia* subgenome of all samples (Figure 2). Similarly, loci heterozygosity remained high for most *D. anglica* and ‘*D. intermedia’* (ID) samples after phasing. This is likely due to partial sequences in the reference transcriptome assemblies, even though we required the reference being present in both parental species. In addition, lineage-specific mutations in the parental and allopolyploid genomes may lead to incomplete phasing. From here on, the subgenome phased to *D. linearis* will be referred to as subgenome L, and the subgenome phased to *D. rotundifolia* will be referred to as subgenome R.

**Figure 3:**
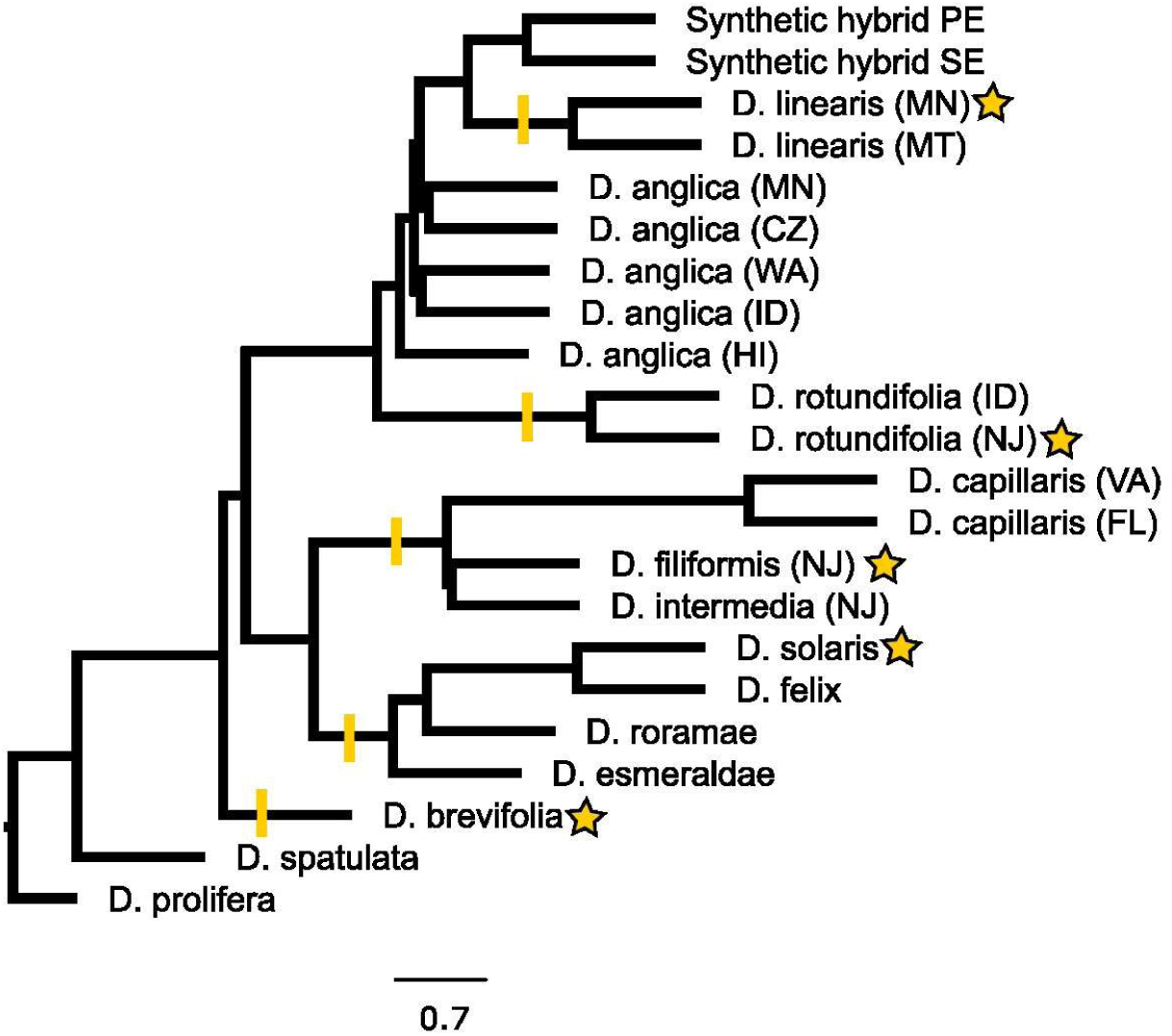
ASTRAL tree estimated from HybPiper assembly without phasing. *Drosera prolifera* was used to root the tree. The scale bar indicates internal branch lengths in coalescent units. All branches have a posterior probability of 1. Terminal branch lengths were artificially chosen. Stars indicate samples used as clade references for phasing, and the yellow lines indicate the clade each clade reference represents.

### Phylogenomic analyses supported the affinity of D. anglica with D. linearis and D. rotundifolia, with D. intermedia nested within separate Eastern North American clades

All phylogenomic analyses, unphased (Figure 3) or phased (Figure 4), supported the affinity of *D. anglica* with *D. linearis* and *D. rotundifolia*—none of the remaining species of sect. *Drosera* were sister to *D. anglica*. From the HybPiper-HybPhaser pipeline, 3057 genes were assembled in every sample or phased subgenome. Phylogenetic analyses using ASTRAL and RAxML recovered very short internal branch lengths (< 0.0005) and discordance between the RAxML and ASTRAL trees among *D. brevifolia, D. linearis, D. rotundifolia,* and the remaining species of the section (Figure 4). All *Drosera* sect. *Drosera* species that occurred exclusively in South America (*D. felix, D. solaris, D. esmeraldae,* and *D. roraimae*) were monophyletic with strong support (supported by 1015/1694 genes each with >50 bootstrap, referred to as informative from now on; Figure 4). Sister to this clade was a clade of eastern North American species, many reaching South America (*D. filiformis, D. intermedia,* and *D. capillaris;* 1543/2128 informative genes; Figure 4). However, species ranging from North to South America were not monophyletic (*D. intermedia, D. capillaris,* and *D. brevifolia*). Across the section, while some closely related species show similar distributions, neither the boreal nor longitudinally distributed species were monophyletic.

**Figure 4.**
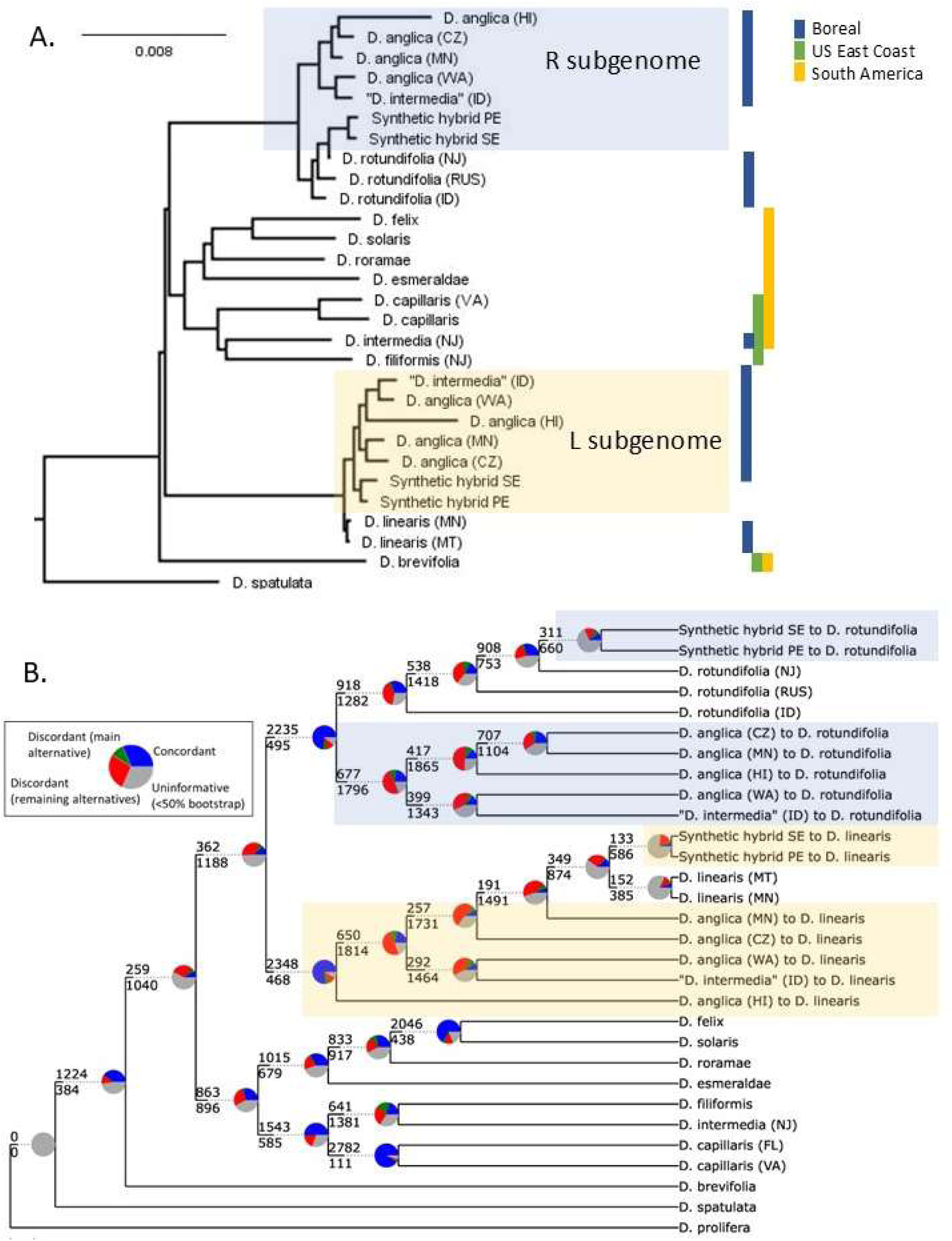
*Drosera* sect. *Drosera* phylogenies estimated from 3569 genes assembled using HybPiper with the subgenomes of *D. anglica* phased using HybPhaser. A. The RAxML phylogram. The scale bar indicates branch length. All nodes had 100% bootstrap support. B. the ASTRAL cladogram with node support from PhyParts.

### The parentage of each D. anglica subgenome is further supported by genetic distance

Using assemblies from the phased reads, both RAxML and ASTRAL recovered sequences from the *D. anglica* R subgenome being monophyletic (677/2473 informative bipartitions) and sister to *D. rotundifolia* + phased subgenomes of the synthetic hybrids. On the other hand, sequences from the L subgenome of *D. anglica* were paraphyletic with *D. linearis* either nested in *D. anglica* (ASTRAL), or monophyletic with *D. linearis* sister to the L subgenome clade (RAxML). Compared to the R subgenome, the L subgenome also had shorter branch lengths, less informative bipartitions, and higher levels of gene tree discordance among populations (Figure 4).

Since the pairwise genetic distance considers gap and base pair comparisons, we further inspected the multiple sequence alignments after phasing. Visual inspection found that removing alignments with an average pairwise genetic distance <98.0% reduced genes with assembly issues. After removing gaps in the alignment and alignments shorter than 1000 bps, 140 genes remained. Overall, the median distance between samples ranged from 0 to 0.023, with the highest distance being between the outgroup *D. spatulata* and *D. anglica* (HI; Appendix S5).

The mean distance between *D. rotundifolia* and *D. linearis* samples was 0.016 (Appendix S5). The mean genetic distance between samples of *D. anglica* subgenome L ranged from 0.0005-0.0010, with a higher value in *D. anglica* (HI) (0.0020-0.0023; Table 2). Similarly, the genetic distance between *D. anglica* subgenome L and *D. linearis* ranged from 0.0005-0.0020, with the highest values from *D. anglica* (HI) (Table 2). On the other hand, there was more variation in the *D. anglica* subgenome R. The median genetic distance between *D. anglica* subgenome R and *D. rotundifolia* ranged from 0.0019-0.0040 (Table 2). The genetic distance between the R subgenomes of *D. anglica* samples ranged from 0.0005 to 0.0028 with the MN-CZ and *’D. intermedia’* (ID) and *D. anglica* (WA) pairs having a genetic distance of only 0.0005 (Table 2).

**Table 2:**
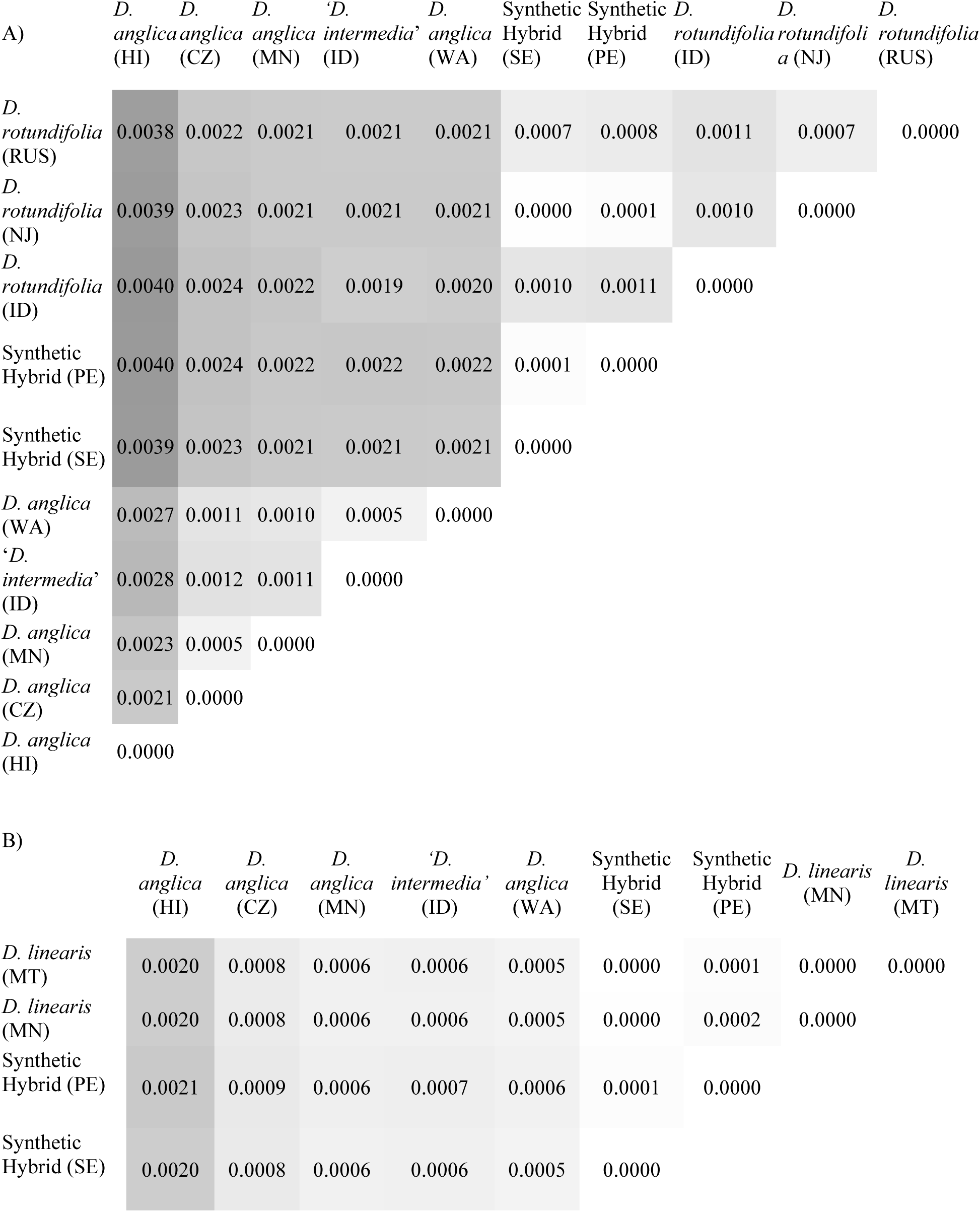
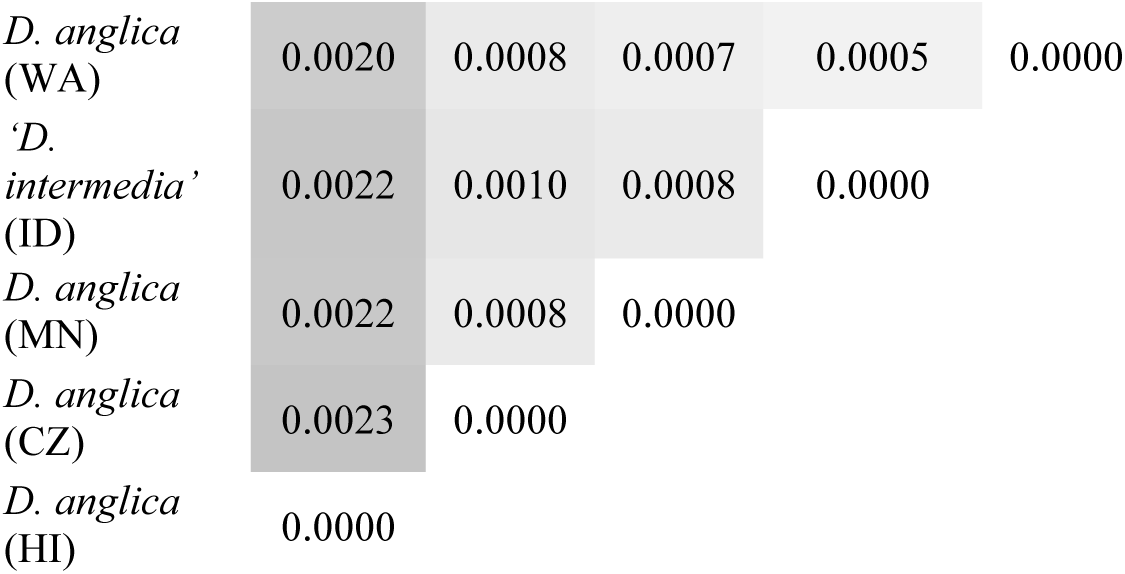
The mean pairwise genetic distance A) between *D. rotundifolia* and *D. anglica* subgenome R and B) between *D. linearis* and *D. anglica* subgenome L from 141 genes after phasing.

| A) | *D. anglica* (HI) | *D. anglica* (CZ) | *D. anglica* (MN) | '*D. intermedia*' (ID) | *D. anglica* (WA) | Synthetic Hybrid (SE) | Synthetic Hybrid (PE) | *D. rotundifolia* (ID) | *D. rotundifolia* (NJ) | *D. rotundifolia* (RUS) |
| --- | --- | --- | --- | --- | --- | --- | --- | --- | --- | --- |
| *D. rotundifolia* (RUS) | 0.0038 | 0.0022 | 0.0021 | 0.0021 | 0.0021 | 0.0007 | 0.0008 | 0.0011 | 0.0007 | 0.0000 |
| *D. rotundifolia* (NJ) | 0.0039 | 0.0023 | 0.0021 | 0.0021 | 0.0021 | 0.0000 | 0.0001 | 0.0010 | 0.0000 |  |
| *D. rotundifolia* (ID) | 0.0040 | 0.0024 | 0.0022 | 0.0019 | 0.0020 | 0.0010 | 0.0011 | 0.0000 |  |  |
| Synthetic Hybrid (PE) | 0.0040 | 0.0024 | 0.0022 | 0.0022 | 0.0022 | 0.0001 | 0.0000 |  |  |  |
| Synthetic Hybrid (SE) | 0.0039 | 0.0023 | 0.0021 | 0.0021 | 0.0021 | 0.0000 |  |  |  |  |
| *D. anglica* (WA) | 0.0027 | 0.0011 | 0.0010 | 0.0005 | 0.0000 |  |  |  |  |  |
| '*D. intermedia*' (ID) | 0.0028 | 0.0012 | 0.0011 | 0.0000 |  |  |  |  |  |  |
| *D. anglica* (MN) | 0.0023 | 0.0005 | 0.0000 |  |  |  |  |  |  |  |
| *D. anglica* (CZ) | 0.0021 | 0.0000 |  |  |  |  |  |  |  |  |
| *D. anglica* (HI) | 0.0000 |  |  |  |  |  |  |  |  |  |

| B) | *D. anglica* (HI) | *D. anglica* (CZ) | *D. anglica* (MN) | '*D. intermedia*' (ID) | *D. anglica* (WA) | Synthetic Hybrid (SE) | Synthetic Hybrid (PE) | *D. linearis* (MN) | *D. linearis* (MT) |
| --- | --- | --- | --- | --- | --- | --- | --- | --- | --- |
| *D. linearis* (MT) | 0.0020 | 0.0008 | 0.0006 | 0.0006 | 0.0005 | 0.0000 | 0.0001 | 0.0000 | 0.0000 |
| *D. linearis* (MN) | 0.0020 | 0.0008 | 0.0006 | 0.0006 | 0.0005 | 0.0000 | 0.0002 | 0.0000 |  |
| Synthetic Hybrid (PE) | 0.0021 | 0.0009 | 0.0006 | 0.0007 | 0.0006 | 0.0001 | 0.0000 |  |  |
| Synthetic Hybrid (SE) | 0.0020 | 0.0008 | 0.0006 | 0.0006 | 0.0005 | 0.0000 |  |  |  |

|  |  |  |  |  |  |
| --- | --- | --- | --- | --- | --- |
| *D. anglica*<br>(WA) | 0.0020 | 0.0008 | 0.0007 | 0.0005 | 0.0000 |
| *D.*<br>*intermedia'*<br>(ID) | 0.0022 | 0.0010 | 0.0008 | 0.0000 |  |
| *D. anglica*<br>(MN) | 0.0022 | 0.0008 | 0.0000 |  |  |
| *D. anglica*<br>(CZ) | 0.0023 | 0.0000 |  |  |  |
| *D. anglica*<br>(HI) | 0.0000 |  |  |  |  |

### rRNA and *rbc*L sequences in D. anglica were nearly identical to D. linearis

*Drosera anglica*’s *rbc*L matched exactly that of *D. linearis* (Figure 5). The maximum read depth for *rbc*L ranged from 32 to 606,594, likely due to different library preparation approaches. Four SNPs distinguished *D. linearis* from *D. rotundifolia*, and all samples of *D. anglica* matched *D. linearis* at all four SNP sites. Four additional sites showed heterozygosity in one of the sample (either RM218 or RM219). The minor variant was found only by duplicated reads sharing the same start, end, and single variant, while the major variant was supported by reads with multiple starts and ends, suggesting that the former was due to a PCR error. Our results suggested that the *D. linearis* lineage represented the maternal parent of *D. anglica*.

**Figure 5.**
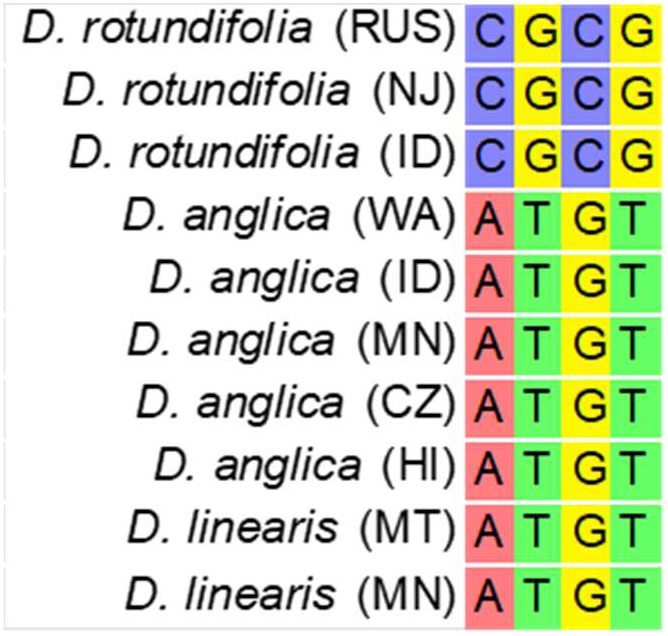
Variable sites in the *rbc*L alignment.

Despite adequate read depth, the ribosomal coding and ITS sequences in both DNA and RNA samples of *D. anglica* were homozygous for SNPs matching *D. linearis*. Read depths reached over 19,000 at regions across the ribosomal RNA locus for all *D. anglica* samples. There were 34 SNP locations, primarily in the ITS region, where *D. linearis* and *D. anglica* were all present and homozygous for one variant, while *D. rotundifolia* was homozygous for the other variant. Five of these SNPs had read depths greater than 2,000 for all *D. anglica* samples. At only one SNP, *D. rotundifolia* and *D. anglica* were homozygous for the same allele, while *D. linearis* was homozygous for a different allele. Two SNPs that were exclusively found in *D. anglica* were homozygous in *D. anglica* (MN), homozygous or heterozygous in *D. anglica* (CZ), and heterozygous in *D. anglica* (HI; Figure 6).

**Figure 6.**
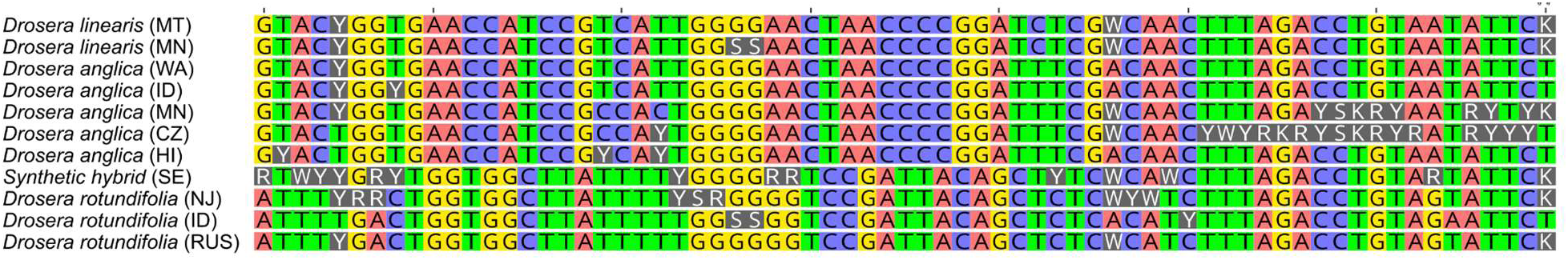
Variable Sites in the ITS alignment. Ambiguity codes: R = A and G, K = G and T, S = G and C, Y = C and T, W = A and T.

### Drosera anglica’s genome has doubled in size compared to most other diploid North American species

We generated genome size estimates for the Washington and Idaho populations of *D. rotundifolia, D. anglica,* and *‘D. intermedia’* (Table 1). Together with previously reported genome sizes, diploid genome sizes for *Drosera* sect. *Drosera* range from 0.6 Gb in *D. spatulata* to 5.9 in *D. tracyi.* North and South American *Drosera* with a chromosome count of 2*n* = 20 had diploid genome sizes mostly between 1.6 Gb and 2.7 Gb except *D. filiformis* and *D. tracyi*, which were 4.9 and 5.9 Gb, respectively. The genome size of *D. anglica* ranged from 4.6 to 4.7 Gb, over twice that of *D. rotundifolia*. Genome size varied between studies and among populations, with our newly generated *D. anglica* ‘WA’ being 75 Mb less than previous estimations of *D. anglica*, and *D. rotundifolia* ‘ID’ was 400 Mb less than previous studies. The *‘D. intermedia’* (ID) population had a genome size more than twice that of *D. intermedia* and similar to, though approximately 660 Mb higher than, the *D. anglica* ‘WA’ population. There are no genome size estimates for *D. linearis*.

## DISCUSSION

By comparing transcriptome and genome sequences in multiple populations across the range of *D. anglica*, we found strong evidence for the origin of *Drosera anglica* from *D. linearis* and *D. rotundifolia.* Further, *D. linearis* represents the maternal lineage and likely contributes to the dominant subgenome, and *D. rotundifolia* represents the paternal lineage if chloroplasts are maternally inherited like most plants. Additionally, we confirmed that the disjunct ‘*D. intermedia’* population in Northern Idaho is *D. anglica* with evidence from both sequences and genome size. Comparing *D. anglica* populations with parental lineages, we found no evidence for multiple independent origins of the European, North American, and Hawaiian populations. More broadly, in *Drosera* sect. *Drosera,* two geographic distributions (boreal and east coast of North America to South America) are not reciprocally monophyletic, suggesting that species diversification in the section may be associated with multiple range shifts. Similarly, leaf length-to-width ratios have shifted at least twice as neither species with narrow long leaves nor short round leaves are monophyletic. Finally, our work presents a cautionary tale that visualization of assemblies and alignments is essential in identifying error and chimeric assemblies and interpreting the data.

### *Drosera linearis* and *D. rotundifolia* are the maternal and paternal parents respectively of *D. anglica*

Sequence assembly of *D. anglica* with Trinity or HybPiper alone resulted in chimeric genes due to the high sequence similarity (98.45%) between the subgenomes. By mapping reads to candidate reference transcriptome assemblies using HybPhaser, we found higher similarity of *D. anglica* subgenomes to *D. rotundifolia* and *D. linearis*, respectively, than to *D. brevifolia, D. filiformis,* or *D. solaris* as representatives of other *D.* sect. *Drosera* lineages. After assembling *D. anglica* reads phased to *D. rotundifolia* and *D. linearis* transcriptome assemblies separately, we found high sequence similarity between *D. anglica* subgenomes and *D. linearis* (99.79–99.95%) and *D. rotundifolia* (99.60–99.80%). Given this high sequence similarity along with the *D. rotundifolia* and *D. linearis* being the only known species in their respective lineages (Mohn et al., 2023), it is unlikely that there is an unsampled species representing either of the parental species. Given that multiple species were tested and compared as alternative phasing references, we were able to exclude the possibility that the resulting subgenome affinity is an artifact of the choice of phasing references. We also performed phasing with HaploSWEEP (Clevenger et al., 2018) and found similar results. However, as HaploSWEEP relies on multiple SNPs occurring on the same read (125 bp), the scarcity of SNPs resulted in very few phased alleles for downstream analyses.

By comparing SNPs in *rbc*L sequences, we determined that the plastid of *D. anglica* originated from *D. linearis*, which likely represents the maternal lineage. While most of our data is based on transcriptomes, our results cannot be explained by RNA editing since *D. rotundifolia* has reduced RNA-editing (Gruzdev et al., 2019), and our genome sequencing results from the Hawaiian population of *D. anglica* recovered an identical *rbc*L sequence compared to those recovered from transcriptomic data. A previous phylogenetic study proposed *D. rotundifolia* being the maternal parent of *D. anglica* as their *rbc*L sequence only differed by three base pairs (Rivadavia et al., 2003). However, Rivadavia et al. (2003) did not sample *D. linearis*, and the similarity reflected the few SNPs between the *rbc*L sequences of *D. rotundifolia* and *D. linearis*.

While the nuclear, bi-parentally inherited ribosomal region is expected to represent both subgenomes following the initial allopolyploidy event, only the *D. linearis* version of the ribosomal region was detected in all our *D. anglica* transcriptome and genome sequence datasets. A similar pattern of gene conversion to a single ribosomal copy (W. Wang et al., 2023) or expression of a single subgenome was observed in both *Gossypium* (Cronn et al., 1999) and *Brassica napus* (Adams et al., 2003). The ribosomal conversion together with a higher number of genes recovered from the L subgenome in both transcriptome and genome datasets suggest that a relatively rare species with a narrow distribution (*D. linearis*) may have contributed the dominant genome to a widespread tetraploid. Alternatively, the difference in genes recovered may be due to undetected phasing issues which were reflected in the L subgenome also having higher allele divergence. Analyses of additional genes in the transcriptomic and genomic data with synteny information are needed to quantify the degree to which the maternal subgenome is dominant genome-wide. The inability to detect both parental lineages in rRNA emphasizes the value of sampling additional nuclear loci in teasing apart subgenomes.

Despite the higher level of genetic similarity between *D. linearis* and *D. anglica,* their homologous chromosomes do not pair properly in hybrids, unlike the homologous chromosomes of *D. rotundifolia* and *D. anglica* (Gervais & Gauthier, 1999; Kondo & Segawa, 1988). The improper pairing of chromosomes suggests chromosome rearrangement events in *D. linearis*, but synteny analysis is needed to determine whether this change is epigenetic or genetic (Fu et al., 2012; Herrera et al., 2008). Gervais and Gauthier (1999) expressed concern that hybridization might dilute and ultimately replace *D. linearis.* The *D. linearis*-specific chromosome rearrangement suggests a potential mechanism that maintains species boundaries between *D. linearis* and the other co-occurring *Drosera* species.

### The northern Idaho population previously referred to as *D. intermedia* is *D. anglica*

All evidence, including genome size, percent of heterozygous loci, allele divergence, and phylogenetic analyses based on both reference-guided and *de novo* assemblies, indicates that the Idaho population of ‘*D. intermedia*’, a critically imperiled species in Idaho (Spoon-Leaved Sundew (*Drosera intermedia*)| *Idaho Fish and Game*, Retrieved: December 20, 2024), is indeed *D. anglica*, a species of not ranked in Idaho (English Sundew (*Drosera Anglica*)*| Idaho Fish and Game*, Retrieved: December 20, 2024). The diploid genome size of ‘*D. intermedia*’ (ID) was about twice that of *D. intermedia* or *D. rotundifolia.* The ‘*D. intermedia*’ (ID) sample had loci heterozygosity and allele divergence similar to the known *D. anglica* populations and much higher than the diploid *D. intermedia*. When mapped to clade representatives, like *D. anglica* and unlike *D. intermedia*, it had a strong affinity to both *D. linearis* and *D. rotundifolia*. In phylogenetic analyses with phased SNPs and haplotypes, ‘*D. intermedia*’ (ID) was nested among the *D. anglica* samples, resulting in the conclusion that this population is a misidentified population of *D. anglica.* From now on, we will refer to it as *D. anglica* (ID).

Despite a strong sequence affinity to *D. anglica,* the genome size of the Idaho population of *D. anglica* was 660 Mb and more than 10% larger than *D. anglica* (WA), greater than the variation observed between fresh and silica-dried samples (Bainard et al., 2011; G. Wang & Yang, 2016) or between *D. anglica* (WA) and previous measurements of *D. anglica* (75 Mb; Veleba et al., 2017). While newly-formed hybrids between *D. anglica* and *D. rotundifolia* or other species are known to occur in wild populations (Mellichamp, 2016), this would not explain the slightly increased genome size. Additionally, the *D. anglica* (ID) genome lacked any evidence of increased similarity to either parent, as might be the case if it had a separate origin or was backcrossed with a parent. Instead, the slightly larger genome in *D. anglica* (ID) may be due to additional repetitive regions or aneupolyploidy.

Confirming this northern Idaho population as *D. anglica* with molecular and genome size data supports Rice’s proposal (2019) based on morphology and habitat. While the leaf shape of *D. anglica* (ID) is similar to *D. intermedia*, the flowering stalks rise vertically from the rosette and its leaves are mostly rising instead of spreading, which are characteristics of *D. anglica* (see Figure 7 for the typical morphology). One potential explanation for the *D. anglica* (ID) population having shorter leaf blades than typical *D. anglica* is its habitat differences from other nearby *D. anglica* populations. While *D. anglica* populations in the region occur on the lakeside edges of floating bogs among *Sphagnum* and buckbean (*Menyanthes trifoliata*) and on floating logs, this population occurs on a sloped fen. Similar leaf-shape plasticity corresponding to habitat differences has been observed in *D. intermedia* (Banaś et al., 2024).

**Figure 7:**
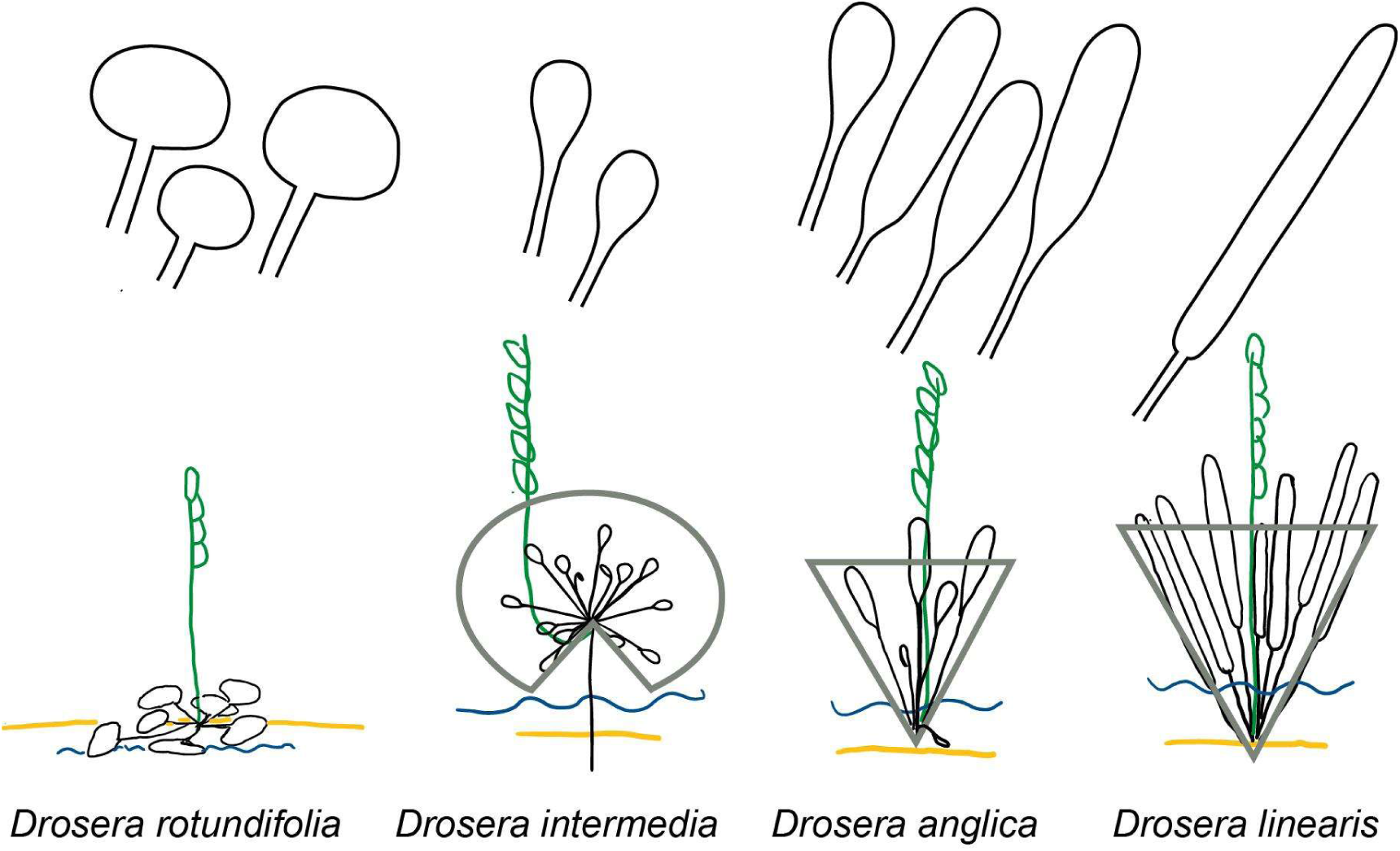
Comparison of leaf shape, angle of the petiole, and shape of the peduncle among *D. anglica,* its parents *D. rotundifolia* and *D. linearis*. We also included *D. intermedia* for comparison as it is often confused with *D. anglica*. The yellow line represents the substrate, and the blue line represents the water level.

This population in northern Idaho and another one in south central Idaho are over 1000 km west of the nearest *D. intermedia* populations, which has made them a conservation priority. With this knowledge of the northern Idaho population being *D. anglica*, the southern Idaho population is likely *D. anglica* as well (Rice, 2019). Our results support removing *D. intermedia* from the list of species and the species of conservation concern in Idaho. More broadly, we observed that *D. intermedia* is often confused with *D. anglica* in herbaria (see Figure 7 concerning distinguishing the two). Our study suggests that additional misidentifications, especially the multiple disjunct populations of *D. intermedia* elsewhere in the world (Figure 1B) may occur due to phenotypic plasticity.

### Hawaiian *D. anglica* originated by long-distance dispersal from the boreal region

The Hawaiian sample of *D. anglica* was deeply nested among *D. anglica* from North America and Europe in the phased data (Figure 4) and showed affinity with remaining *D. anglica* samples in unphased data (Figure 3). This Hawaiian *D. anglica* sample had higher allele divergences and terminal branch lengths than the remaining *D. anglica* samples. This may be partially due to using DNA instead of RNA for the Hawaiian *D. anglica* sample. We are likely to recover a more balanced and complete representation of alleles and homeologs from genomic than from transcriptomic sequencing. To ensure that the higher sequence divergence was not an artifact from intron or chimeric assemblies from stitching together exons, before and after phasing we also reanalyzed the *D. anglica* (HI) genome sequence data without stitching contigs in HybPiper, but the result was only a slightly decreased allele divergence. The higher genetic diversity found in the Hawaiian sample is unlikely to be due to a higher ploidy level, as a previous study found a chromosome count of 2n=40 for a Hawaiian population of *D. anglica* (Hoshi & Kondo, 1998). As *D. anglica* is native to only the island of Kauài, restricted to the montane wetlands on the island (Mellichamp, 2016), the chromosome count likely came from the same or nearby population to our DNA sample. More likely, the higher genetic diversity and longer terminal branch lengths are due to the severe bottleneck experienced by long-distance dispersal to the remote oceanic island, a longer growth season with the loss of dormancy (Mellichamp, 2016), and other adaptations to the tropical mountain forest habitat of the Hawaiian population.

### No evidence supports multiple origins of *D. anglica*

Given the circumboreal distribution of both *D. anglica* and its paternal parent *D. rotundifolia*, we sampled populations of both species from North America and Europe to test whether *D. anglica* originated multiple times. However, we did not find evidence supporting multiple independent origins of *D. anglica*. While *D. anglica* subgenome L was paraphyletic in the ASTRAL tree estimated from the phased data, the topology had little support (gray and red in Figure 4B). In fact the genetic distance between each subgenome and its parental lineage ranged from 0.001 to 0.004, which equates to a few mutations at most per gene. Regions with higher evolutionary rates may be more informative for further testing multiple origins. The higher genetic diversity in the R subgenome among *D. anglica* populations may still come from multiple origins, larger effective population sizes in the ancestral *D. rotundifolia* compared to the *D. linearis* genomes, or higher levels of purifying selection in the L subgenome given its genomic dominance.

The larger population size and widespread distribution of *Drosera rotundifolia* throughout the Pleistocene has been supported by multiple lines of evidence. Pleistocene fossils of *D. rotundifolia* have been found in Canada (Penhallow, 1890, 1896). In addition, Korean populations of *D. rotundifolia* showed low within-population diversity but high between-population divergence, suggestive of micro-refugia during the last glacial maximum (Chung et al., 2013). On the other hand, *D. linearis* is more restricted in its geography and habitat, and the flarks where it occurs expand and contract more rapidly with climate fluctuations (Kolari et al., 2022). This may explain the lower genetic diversity in *D. linearis* than in *D. rotundifolia*.

*Drosera anglica*, the allopolyploid hybrid, occurs in an intermediate habitat between bogs and fen flarks that is more abundant than the flarks of *D. linearis* and likely experienced less fluctuation of population sizes than *D. linearis*.

### *Drosera anglica* is intermediate between both parents in morphology and microhabitat

The leaf shape, angle of the petiole, and shape of the peduncle distinguish *D. anglica* from its parental species and the frequently confused *D. intermedia* (Figure 7). *Drosera rotundifolia*’s leaf blade is wider than long, the leaves are generally flat against the ground or slightly raised, and the peduncle rises directly from the center of the plant. *Drosera anglica* and *D. linearis* leaves are generally raised, although older leaves may spread some. Both species have peduncles originating vertically from the plant like *D. rotundifolia* but unlike *D. intermedia,* which has a peduncle that emerges close to horizontally before it curves upward. *Drosera linearis*’s leaf blades are linear, as its name suggests, with the two sides of the leaf being parallel and the ends stopping abruptly instead of tapering. *Drosera anglica*’s leaf shape is quite variable, ranging from oblong to linear-spatulate (Lowrie et al., 2017; Mellichamp, 2016). In mature *D. intermedia* plants, the leaves are spread out evenly and may be reflexed when the plant has a stem. Like the shorter of *D. anglica*’s leaves, they tend to be spatulate (Lowrie et al., 2017).

The differences in leaf shape and orientation among *D. rotundifolia, D. anglica, D. linearis,* and *D. intermedia* correspond to their different microhabitats. At the two extremes, *D. rotundifolia* occurs in drier conditions on sphagnum hummocks, and *D. linearis* occurs in the standing water of fen flarks, naturally occurring swales on a patterned peatland. *Drosera intermedia* can occur standing in water or on drier peat and tends to be found in drier habitats than *D. anglica. Drosera anglica* is often standing in water on the edges and ridges of peatlands and on the edges of floating bogs (Banaś et al., 2023).

When species co-occur, putative hybrids are occasionally observed with intermediate leaf shapes and habitats (Gervais & Gauthier, 1999; Lowrie et al., 2017; Wood, Jr., 1955). For example, in northern Minnesota we found a plant growing with *D. rotundifolia* on a dry hummock and with *D. linearis* nearby. While the habitat suggested *D. rotundifolia,* it had spatulate-shaped leaves that were more upright. It had a genome size similar to that of *D. rotundifolia.* This plant produced well-filled seeds, although we have not been able to germinate the seeds after testing a small batch of eight.

### *Drosera* sect. *Drosera* has undergone multiple shifts in leaf shape and range

Although leaf shape and position are generally reliable in distinguishing *D. anglica* vs. close relatives, each type of leaf shape is non-monophyletic across *D.* sect. *Drosera.* Neither long skinny leaves (*D. filiformis*, *D. linearis*) nor round leaves (*D. rotundifolia*, *D. brevifolia*, *D. capillaris*) and neither erect leaves (*D. filiformis*, *D. linearis*, *D. intermedia*, *D. roraimae*) nor prostrate leaves (*D. brevifolia*, *D. capillaris, D. esmeraldae,* and often *D. rotundifolia*) are monophyletic (Lowrie et al., 2017; Mellichamp, 2016). Leaf shape and position are important for prey capture as together they position the leaf blade with sticky traps above water or surrounding plants (Fleischmann et al., 2018). Likewise, within the section, the two major distributions— boreal, and Eastern North America to South America—are non-monophyletic, indicating that range shifts have occurred multiple times. Of the 15 sections of *Drosera, D.* sect. *Drosera* occupies the broadest range in climate and geography. The broad climatic adaptation and geographic distribution is likely achieved via repeated shifts in strategies in prey capture and in adaptations to macro- and micro-habitats.

### Data visualization at multiple stages is important in analyzing large datasets

Our work presents a cautionary tale highlighting the importance of data visualization in multiple steps of data analyses to identify unexpected issues. These issues included sequence processing errors, violated assembly assumptions, and issues in the quality of reference genome annotations.

Visualizing read mapping in IGV resulted in the detection of a sequence processing error. After cleaning reads and assembling genes, we observed that all single-end read samples from the same sequencing batch had an increased number of variants on the 3’ end. In addition to residual adaptors, we observed a single ‘T’ on the 3’ end of many reads. When enough reads with a terminal ‘T’ ended at the same place, this resulted in an SNP being called erroneously and over-estimating divergence.

HybPiper was designed to assemble genomic reads to coding sequence targets. Therefore, when ends of reads do not map to the target, they are assumed to be introns and are trimmed.

When we adopted HybPiper to assemble transcriptomic reads to coding sequence targets from genome annotation, we observed assemblies that appeared to have even coverage across the gene, but the reads had been trimmed, and no reads were bridging two adjacent regions. This is likely due to either alternative splicing or chimeric sequences in the reference. However, we observed other regions which were quite likely chimeric sequences in the reference genome because when we visualized the reads mapped to the *D. spatulata* reference, we observed that some genes had one region with read depths in the thousands while other regions had much lower read depths. Often, no reads spanned these two regions, suggesting the presence of chimeric genes in the *D. spatulata* genome annotation.

By visualizing our assemblies, we identified issues with read processing errors in three samples, issues with chimeras in the reference transcripts from a genome assembly, and issues with applying a target enrichment pipeline to transcriptomic data and were able to take corrective measures. However, these issues could be easily overlooked without the visualization of assemblies and alignments and could have propagated in our results by overestimating the divergence of sequences.

## Conclusion

Both reference-guided and *de novo*-based read assembly methods supported *D. rotundifolia* and *D. linearis* as the paternal and maternal lineages of *D. anglica*, respectively, with evidence of the maternal subgenome being dominant. We found no evidence for multiple origins of *D. anglica*. The Hawaiian *D. anglica* originated through long-distance dispersal from the boreal region Future work should further explore subgenomic dominance and population structure in *D. anglica,* including *D. anglica* from Hawaii, and evaluate the presence of a chromosomal rearrangement in *D. linearis*.

Our work highlighted the importance of careful examination of phylogenomic data, including testing for cross-contamination, and visual inspection of read mapping, SNP patterns, alignments, and individual gene trees. Simply running through standard phylogenomic pipelines would likely omit cryptic reticulation events that are of ecological, evolutionary, and conservation importance.

## ACKNOWLEDGEMENTS

The authors would like to thank Bruce Mohn, Caleb Mohn, Darren Loomis, Ezra Mohn, Kristen Bednarczyk, Stephen Mohn, and Jason Husveth for field assistance; Andrea Pipp, Barry Rice, Chel Anderson, Chris Ludwig, Ethan Perry, John Hayden, John Townsend, Michael Lee, Nicholas Severson, Shannon Kimball, Tim Whitfeld, and Welby Smith for information on accessing plant populations. The authors would also like to thank the Idaho Panhandle National Forest and the Helena National Forest; the Minnesota Department of Natural Resources; New Jersey Bureau of Land Management, Division of Fish and Wildlife; and Virginia Department of Natural History for granting access to study sites. The authors would also like to thank Alex Eilts, Lisa Philander, and Jared Rubinstein at the University of Minnesota College of Biological Sciences Conservatory for assistance in cultivating the plants and Clifford Morden for providing DNA samples. The authors would like to thank Yinyin Huang and Zack Radford for their lab assistance, Diego Morales-Briones and Aaron Lee for help with data analysis, and Alan Whittemore, Marlene Hahn, Lindsey Worcester, Alison Branz, and Alex Krupa for their feedback on the manuscript. Sequencing was performed at the University of Minnesota Genomics Center Facility (Minneapolis, MN) and Novogene Corporation, Inc.

## AUTHOR CONTRIBUTIONS

RM designed the study and led sample collection, lab work, analysis, and writing. YY assisted with study design, sample collection, analytical approaches, and writing.

## DATA AVAILABILITY STATEMENT

Raw transcriptomic and genomic reads for newly generated samples will be made available in the NCBI SRA PRJNA1198097 before publication. Assembled sequences, multiple sequence alignments, and trees will be made available through Dryad (https://doi.org/10.5061/dryad.7m0cfxq5j). Additional supporting information may be found online in the Supporting Information section at the end of the article

## APPENDICES

**Appendix S1:** The modified PureLink RNA extraction and DNase protocols.

**Appendix S2:** Genome sizes estimation and newly sequenced sample information. 1) Sample information and genome size estimations, 2) Sequencing and assembly information, 3) HybPhaser association of samples with clade references, and 4) Allele divergence, % locus heterozygosity, and number of loci recovered, and

**Appendix S3:** Photo vouchers.

**Appendix S4:** Ks plots with Ks values 0 to 0.5.

**Appendix S5:** Genetic distances between all samples. 1) Median and 2) mean pairwise genetic distance between samples.

