## Appendix S1 for "Origin of subgenomes in the circumboreal allopolyploid carnivorous plant *Drosera anglica* (Droseraceae)"

### PureLink RNA Extraction

This is modified from (Jordon-Thaden et al., 2015; Yang et al., 2017).

#### Preparing Equipment

1. Get liquid nitrogen and crushed dry ice.
2. Use liquid nitrogen to prepare cool rack in Styrofoam box.
3. Pre-cool the refrigerated centrifuge to 4°C
4. Reagents: PureLink reagent , 5 M NaCl, 75% ethanol, Isopropanol,
5. Optional Reagents: Plant RNA Isolation Aid
6. Prepare: 1.5% agarose gel

#### Preparing space

1. Wipe the workspace with 70% alcohol followed by RNaseZap.
2. Label on the side of tube, tap down the beads. Place tubes on rack in liquid nitrogen to let them chill.

#### Collecting Sample

1. Clean tweezers with 70% ethanol and RNaseZap. Dip the tweezers in liquid nitrogen to chill. Remove < 0.1 g tissue from the bottle (approximately size of a punch hole; can skip weighing to avoid thawing). Record tissue types on lab notebook.
2. Clean and prepare the tweezers by spraying with 70% ethanol and RNaseZap as in step 5, and dip in liquid nitrogen to chill before proceeding to the next sample. Add liquid nitrogen to the styrofoam box when it’s low.

#### Grinding Sample

1. Tape the cryogenic adapter except the number of tubes you are running (with half on each side) so that the dry ice won’t fly out while shaking. Transfer < 0.5 g crushed dry ice to the FastPrep adapter. Use small pieces so that it’s easier to balance.
    a. Note: Running 2-4 samples is a lot faster and more manageable.
2. Grind frozen tissue in the FastPrep-24 on Adapter: CoolPrep and 4 m/s for 40 seconds (program manually). After finishing, immediately move lysing matrices back to the rack in dry ice to avoid thawing. Tap down the bead gently on bench.
3. Add more dry ice to the adapter and grind for another 40s. Put lysing matrix back onto the rack in liquid nitrogen. The tissue should be in very fine powder. If not, repeat for a third round of grinding.

    a. Do not grind for more than 3 rounds as tubes may break.

**RNA extraction**

Perform all steps in the hood. The samples need to be **kept frozen** until PureLink reagent is added.

1. Aliquot PureLink reagent into a 50 mL falcon tube. Volume needed = # samples * 500mL + 300 mL. (For example, 6.3 mL for 12 samples)
2. Estimate the weight of each sample. Calculate the amount of Plant RNA Isolation Aid (PRIA) using the following formula 1mL of PRIA/ gram of material. Add 30 extra microliters and aliquot enough Plant RNA Isolation Aid into a microcentrofuge tube:
    1. Small: 0.02-0.03 grams = 30 uL
    2. Medium-Small: 0.05 g = 50 uL
    3. Medium-Large: 0.07-0.08 g = 80 uL
    4. Very Large: 0.1 g = 100 uL

    (For example, 1 small + 1 medium + 2 medium large + 30 mL= 270 uL)

3. Add 500ul PureLink reagent to the frozen, ground plant tissue
4. Immediately, Add 1mL of Plant RNA Isolation Aid/ gram of material to the lysate following estimations above.
5. Immediately, tighten the lid and vortex and occasionally invert the tube until the sample is thoroughly thawed and resuspended with no clumps at the bottom of the tube. Put the tube in a clean, **room-temperature** rack.
    1. In the grinding process tissue can be packed into the lid, so inverting helps stabilize the RNA in the lid as well.
6. Incubate the tube horizontally for 5 minutes at **room temperature**.
7. Meanwhile label a new set of microcentrifuge tubes and add 100uL of 5 M NaCl to each empty new 1.5 mL tube.
8. Centrifuge the sample tubes at 12,000 × g for 2 minutes at **room temperature**.[rcf=xg]
9. Use 200 µL tips to transfer the supernatant to the new tubes with 5 M NaCl. (often slightly more than 400 µL). After transfer, pipette up and down **gently** to mix the supernatant with NaCl. Do not vortex to mix.
10. If the extract is very sticky, add an extra 100 µL or RNase free water to the tube.
11. Add 300uL chloroform to each sample, and mix thoroughly by vortex.
12. Centrifuge the sample at 12,000 × g for 10 minutes at **4°C** to separate the phases.
13. Label a new set of tubes and add 300 uL of chloroform. (this second round of chloroform, especially when the middle layer is pretty thick is helpful for cleaning)
14. While avoid disturbing the middle layer, transfer the upper, aqueous phase using 200 µL tips to the new tubes with chloroform. (This should be about 400 uL, but it is better to move less than disrupt the phase separation.) Mix new tubes thoroughly by vortex.
15. Centrifuge the sample at 12,000 × g for 10 minutes at **4°C** to separate the phases.

16. Label a new set of tubes with sample ID on top, and date and tube number on the side of the tube. Add 350 µL of isopropyl alcohol to each empty tube.
17. Set the p200 pipette to 175µL. Transfer 350µL of the upper, aqueous phase using 200 µL tips into the new tubes with isopropyl alcohol. Make sure not to disturb the middle layer. Pipette to mix and let stand at **room temperature** for 10 minutes.
    1. It is better not to disturb the middle layer than to get all the upper aqueous layer.
18. Centrifuge the sample tubes at 12,000 × g for 10 minutes at **4°C**.
19. Decant/ Pipette off the supernatant, do not disturb the pellet. Add 1 mL of 75% ethanol to the pellet.
    1. Decanting can leave some residue on the outside of the tube, so I prefer to pipette off.
20. Centrifuge at 12,000 × g for 2 minute at **room temperature**. Decant/Pipette off the supernatant without losing the pellet.
21. Briefly centrifuge to collect the residual liquid and remove it with a 20-µL pipette. Leave the tube open to air dry for 15–30 min.
    1. The pellet may turn clear/translucent as it dries.
22. Add 40-60 µL RNase-free water to the RNA pellet. Pipet the liquid up and down over the pellet to resuspend the RNA. It is OK if the solution is still cloudy or pigmented after mixing. It will be cleaned up at the DNase step.
    1. In Drosera it is hard to get multiple tubes from the same plant, so diluting with more water means that it is less consumed by QC steps, so often I go with 50 for a sample with plenty of tissue but no back-up tube.
23. Add 1 µL of RiboLock per 40 µL of water to the sample, and mix.
24. Place sample on ice to slow possible degradation.
25. Visualize of 3 µL of RNA on the 1.5% agarose gel. It's OK to use a DNA ladder. Purified RNA can be kept at **4°C** for a day or two, or at **-80°C** for long-term storage. Alternatively, proceed to DNase step immediately.
26. Pour waste into the waste container. Wash the room temperature racks with tap water. Pour waste liquid into the extraction waste collection bottle in the hood. Discard tips and tubes in the sealed ziplock bag to the hazardous waste bucket. Allow leftover dry ice and liquid nitrogen to evaporate on lab bench and wash the containers and rack sitting in liquid nitrogen the next day.

DNase

This is modified from (Jordon-Thaden et al., 2015; Yang et al., 2017).

**Equipment**

In addition to the equipment required for the PureLink RNA extraction protocol, you will need:

- Dry heating block that holds 1.5 μL tubes (preferred) or an incubator.
- Invitrogen™ TURBO DNA-free™ Kit (Thermo Fisher Scientific, Waltham, Massachusetts, USA), stored in -20°C freezer.
- Agilent 2100 Bioanalyzer and Agilent RNA 6000 Nano Kit (Agilent, Santa Clara, California, USA). Sequencing cores also usually provide Bioanalyzer service

**Prepare Equipment and Samples**

1. Take the DNase buffer out of the -20°C freezer to thaw at room temperature.
2. Turn on the dry heater or incubator to preheat to 37°C.
3. If you have two tubes per sample with similar quality, combine them to increase yield and diversity of genes (total of ~70 μL). Vortex the DNase buffer and spin it down briefly. Add 0.1 volume of 10X Turbo DNase buffer to each tube. For 37 μL of RNA add 3.7μL buffer.

**DNase**

4. Add 1 μL of DNase from the TURBO DNA-free Kit to the RNA. Watch closely to make sure the 1 μL is indeed transferred into the RNA solution. Vortex briefly to mix.
5. Incubate at 37°C for 30 minutes. While waiting, label new 1.5 mL storage tubes with the collection number on top, and the date and tube number of extraction on the side.
6. Add vortexed DNase Inactivation Reagent in the TURBO DNA-free Kit (typically 0.1 volume; 3.7 μL for 37 μL of starting RNA) and mix by vortex briefly. Incubate at room temperature for 5 minutes, vortex occasionally.
7. Centrifuge at 10,000 x g for 2 min and transfer 37 μL (or original volume) of supernatant to the new, pre-labeled storage tubes, and aliquot 4 μL for Bioanalyzer. Place cleaned RNA in 4°C if library prep will be performed in the following day or two. Otherwise store at -80°C.
    a. Disturbing the pellet can cause degradation of the mRNA, so it is better to only pick up 37 μL (or a few microliters less than are in your tube) than to get the last little bit.
8. Run the cleaned RNA on a Bioanalyzer using the Agilent RNA 6000 Nano Kit chips.
    a. Be sure to check the Bioanalyzer results by eye as the peaks may be (are often) mislabeled so a low RIN number may actually be fine.
    b. Mucilaginous tissue can give distorted Bioanalyzer traces but library prep might be OK. Repeat the DNase digestion a second time if a high molecular

weight DNA band shows up. Chloroplast rRNA gives additional bands and can appear as a smear on an agarose gel but will be distinguishable on Bioanalyzer trace.

**Other Notes:**

1. We tried other kits including Qiagen RNeasy, RNAqueous total RNA, but no RNA was obtained suggesting that the columns were clogged in the process.
2. Purelink becomes less effective over time, especially if over 1.5 years old. If extractions are becoming degraded, consider replacing the Purelink reagent.
