## Supplementary figures and images for "Origin of subgenomes in the circumboreal allopolyploid carnivorous plant *Drosera anglica* (Droseraceae)"

### Appendix S4

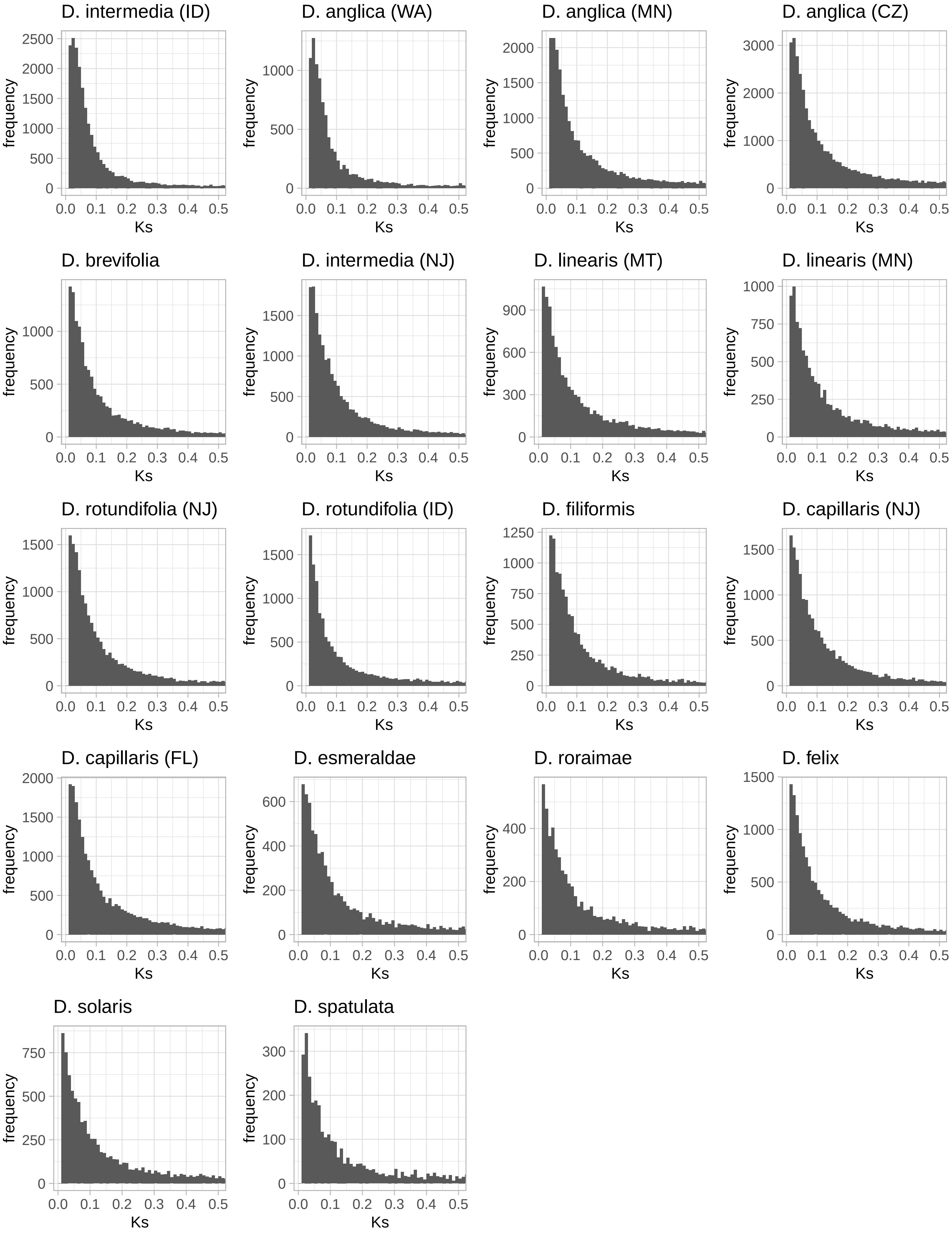

### D_anglica_CZ_RM298.jpg

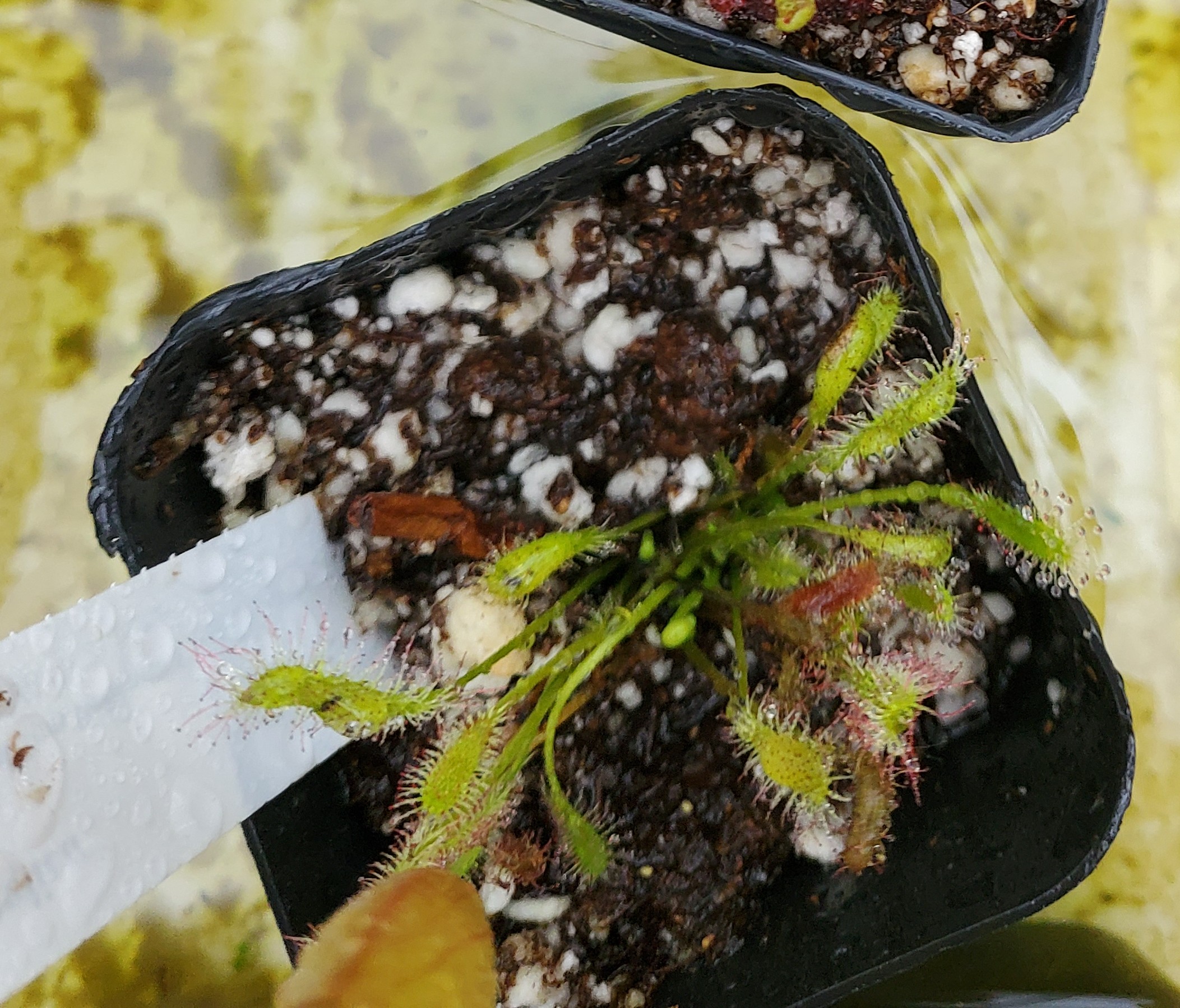

### D_capillaris_FL_RM240_1.JPG

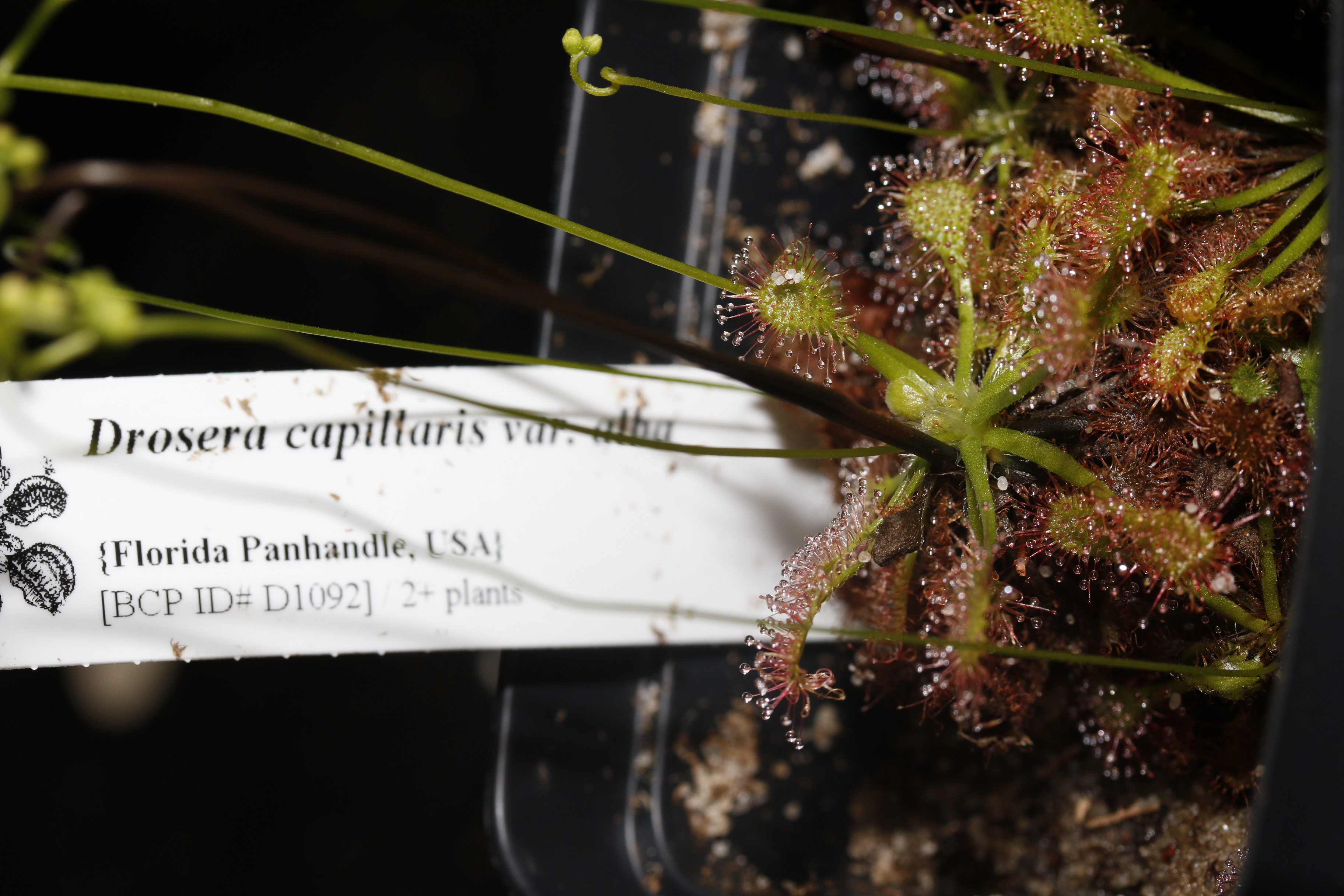

### D_esmeraldae_RM241_2.JPG

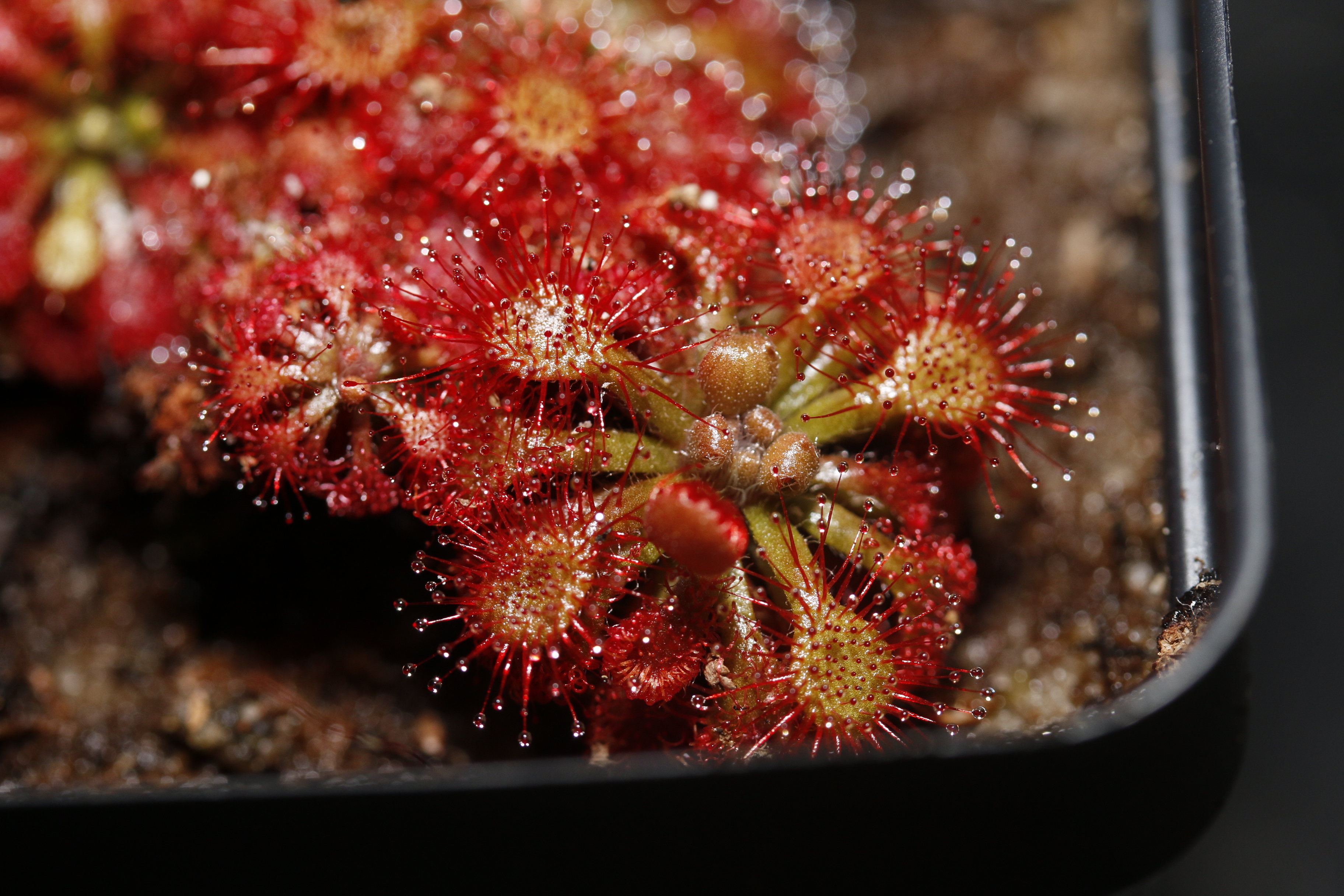

### D_esmeraldae_RM241_3.jpg

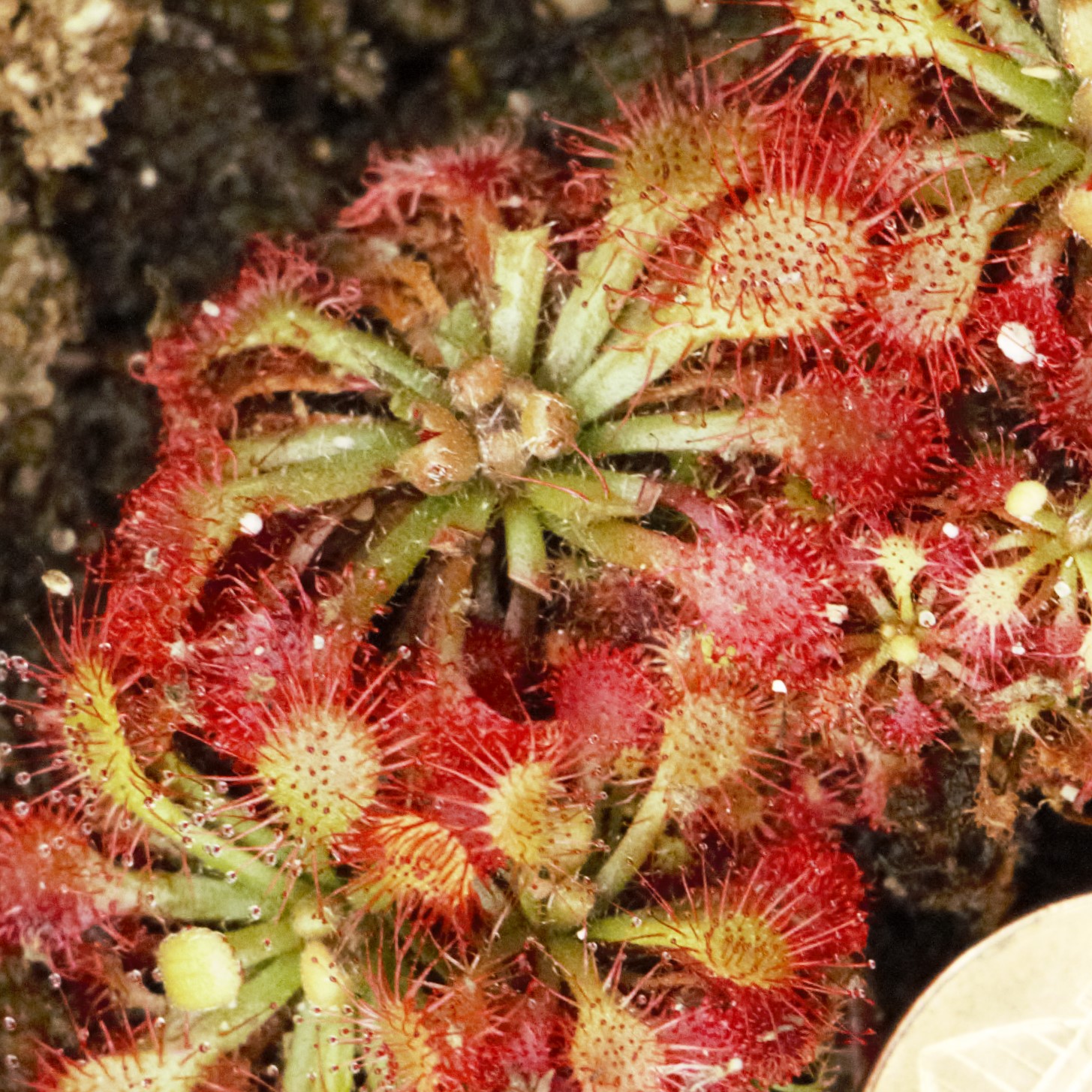

### D_felix_RM245_1.JPG

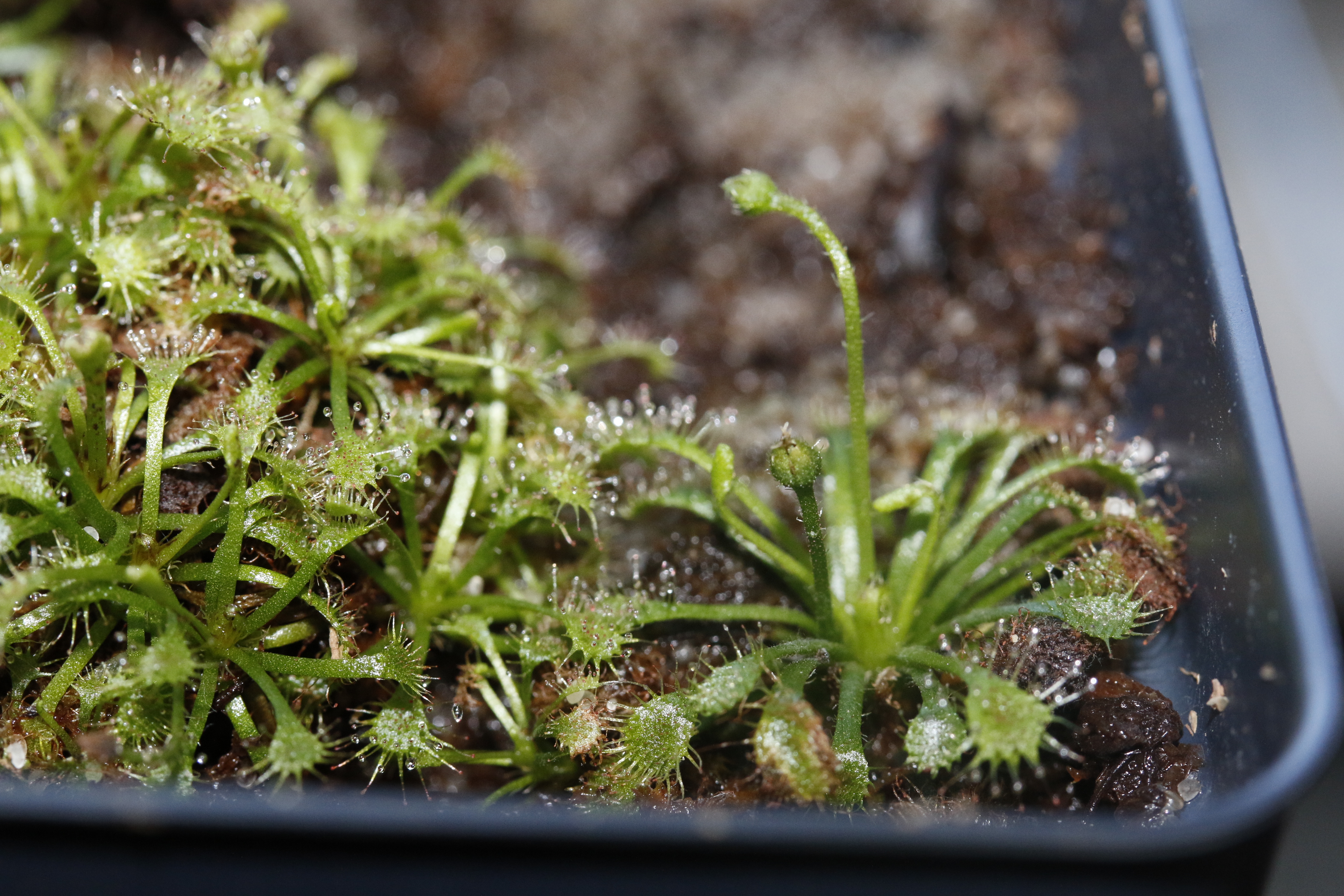

### D_roraimae_RM242_1.JPG

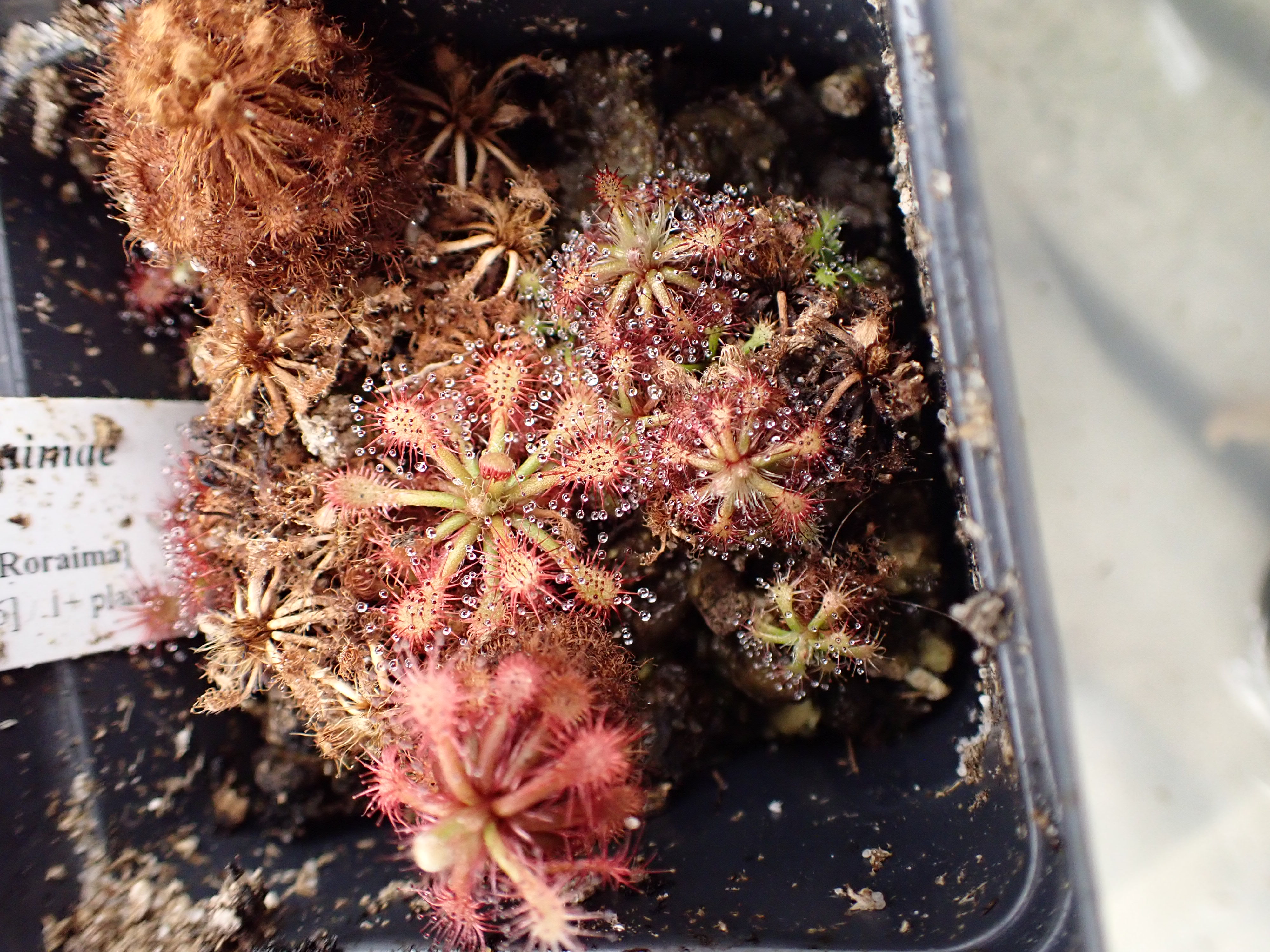

### D_roraimae_RM242_2.JPG

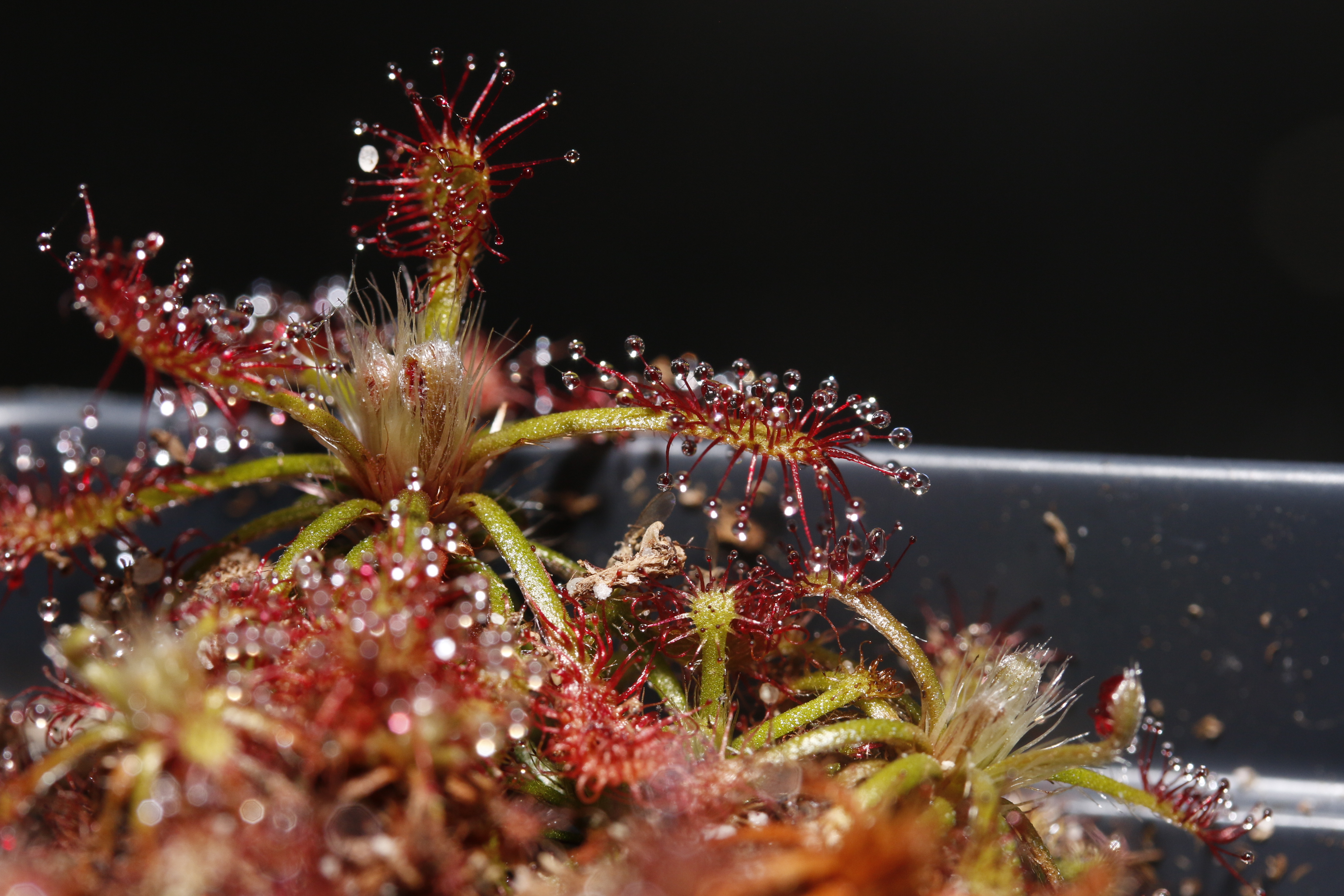

### D_solaris_RM237_1.JPG

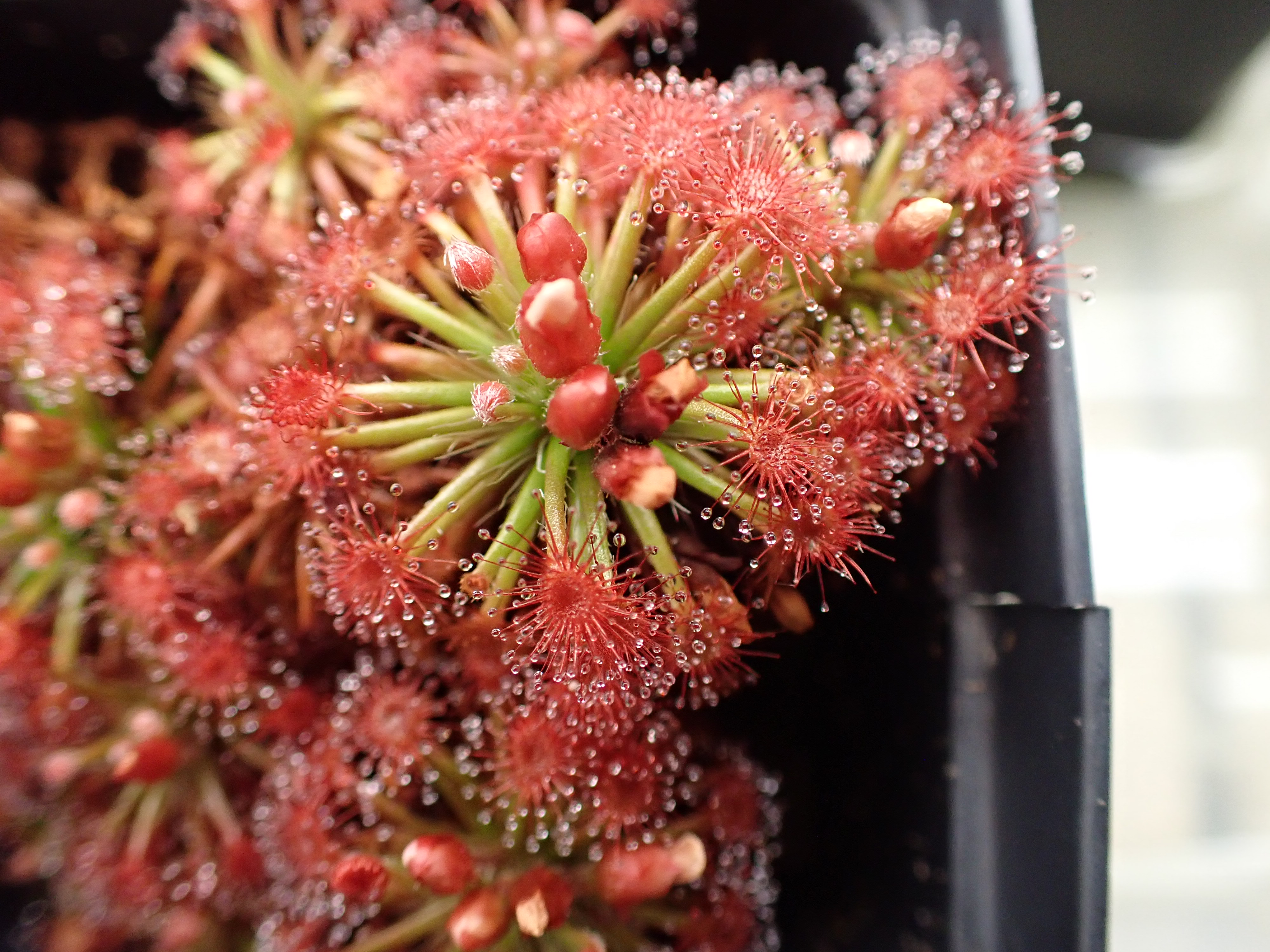

### D_solaris_RM237_2.JPG

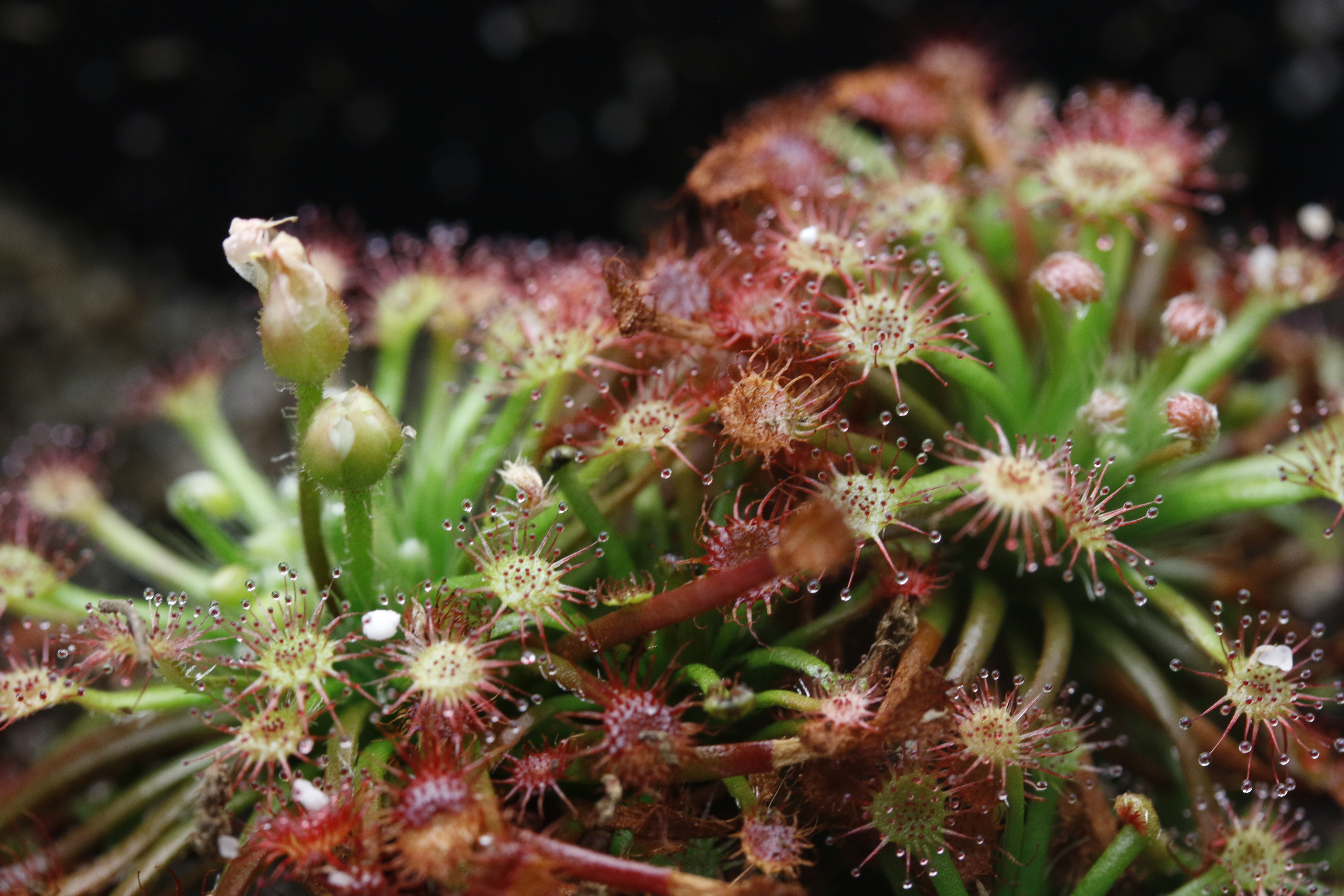
